# 3D Bioprinted Cell-laden GrooveNeuroTube: A Multifunctional Platform for *Ex Vivo* Neural Cell Migration and Growth Studies

**DOI:** 10.1101/2025.02.19.639097

**Authors:** Jagoda Litowczenko, Yannick Richter, Hawrez Ismael, Łukasz Popenda, Adam Ostrowski, Katarzyna Fiedorowicz, Jose Carlos Rodrigez Cabello, Jacek K. Wychowaniec, Krzysztof Tadyszak

## Abstract

Extensive peripheral nerve injuries often lead to the loss of neurological function due to slow regeneration and limited recovery over large gaps. Current clinical interventions, such as nerve guidance conduits (NGCs), face challenges in creating biomimetic microenvironments that effectively support nerve repair. The developed ***GrooveNeuroTube*** is composed of hyaluronic acid methacrylate and gelatin methacrylate hydrogel, incorporating active agents (growth factors and antibacterial agents) encapsulated within an NGC conduit made of 3D-printed PCL grid fibers. In vitro studies showed that ***GrooveNeuroTube*** significantly promoted migration of dorsal root ganglion (DRG) neuronal cells, 3D bioprinted at the far ends of the conduit to imitate a proximal nerve injury as a novel *ex vivo* model. A long-term culture of up to 60 days was employed to better mimic *in vivo* conditions. This model tested the effects of pulsed electromagnetic field (PEMF) stimulation on neural tissue development. After 60 days, ***GrooveNeuroTube*** showed a 32% cell migration increase compared to the growth-factor-group and 105% compared to the no-growth-factor condition. These results confirm that the ***GrooveNeuroTube*** system can effectively support sustained neural cell migration and maturation over extended periods, proving a new technology for testing peripheral nerve injury *ex vivo*.

**Graphical abstract:** 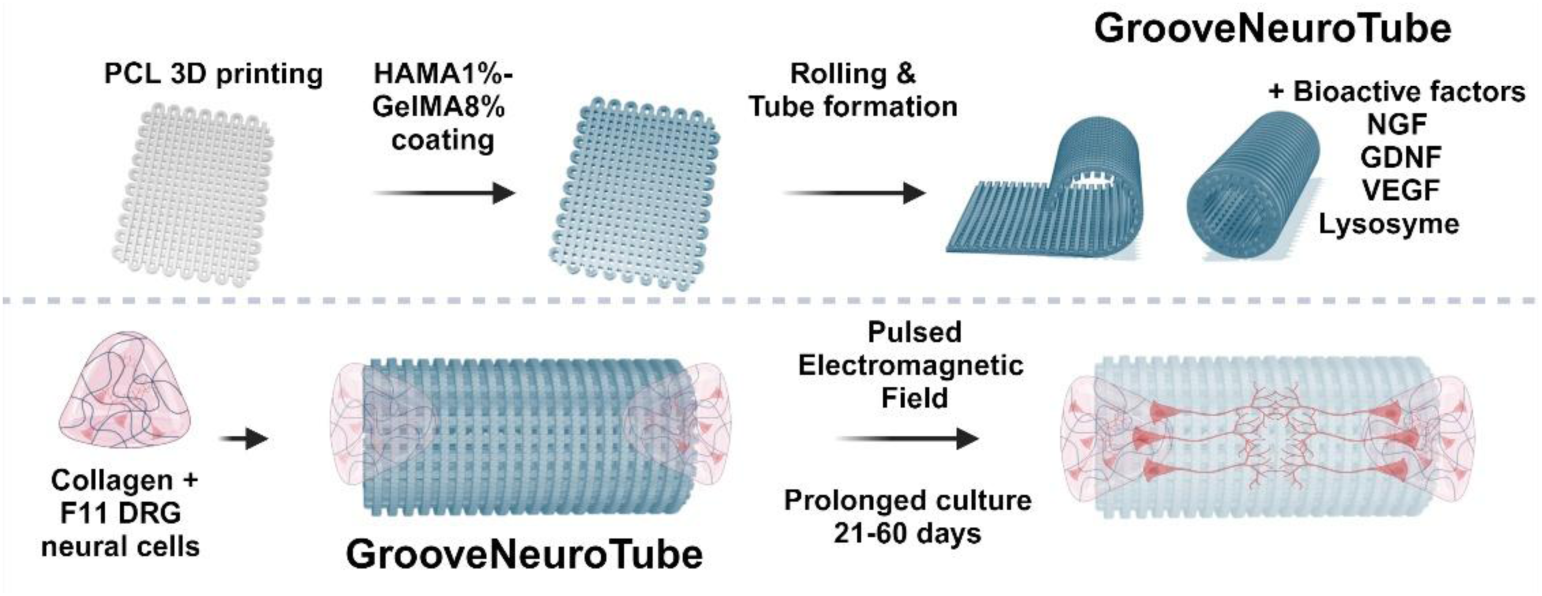

The graphical abstract was created with BioRender.com.

## 1. Introduction

Extensive peripheral nerve injuries (PNI) represent a significant clinical challenge, often leading to substantial functional deficits and prolonged recovery due to the limited capacity for repair.[1, 2] Although the peripheral nervous system (PNS) possesses some regenerative abilities, clinical outcomes following PNI remain suboptimal, highlighting the necessity for advanced therapeutic strategies. Severe PNIs, particularly those with segmental nerve defects exceeding 4 cm, present additional complexities as direct end-to-end repair generates detrimental tension, hindering recovery.[3] In such cases, nerve grafts or conduits designed to bridge the gap have become a cornerstone and necessity of nerve repair. Autologous nerve grafting, considered the gold standard, is still widely employed but is associated with several drawbacks, including donor site morbidity, size mismatches, and the limited availability of donor nerves.[4] As an alternative, nerve guidance conduits (NGCs) constructed from biomaterials have gained significant attention.[1] These conduits can be made from natural materials like collagen, which offer biocompatibility,[5] or synthetic materials like polycaprolactone (PCL), which allow tuneable mechanical and degradation properties.[6] Furthermore, integrating advanced technologies, such as topographical patterning or the addition of bioactive agents, can enhance axonal alignment and nerve regeneration.[4, 6, 7] Topographical patterns can be especially suitable in neural tissue engineering to produce neural conduits to enhance axonal growth and nerve regeneration. Axon outgrowth and pathfinding are crucial for re-establishing neuronal circuitry in re-wiring damage from neuronal injury.[8–11] Biofabrication technologies enable rapidly developing more complex NGCs with custom designs and hierarchical arrangements that can leverage individual requirements for future personalized approaches.[12]

Hyaluronic acid (HA) is a biocompatible and biodegradable glycosaminoglycan found in the extracellular matrix [13] known for its moisture retention, immunomodulatory properties,[14] and gel-like characteristics that support tissue regeneration.[15–18] It plays a critical role in cell adhesion, proliferation, and wound healing, making it valuable for neural tissue engineering (NTE).[19] HA-based hydrogels have been shown to promote neuronal outgrowth and differentiation, offering the potential to treat neurodegenerative disorders.[19]

Studies have demonstrated that HA hydrogels enhance neural cell survival, influence neural patterning, and improve neurological function by releasing neurotrophic factors, making them promising for CNS injury treatment.[19] Soft HA hydrogels, in particular, significantly enhance neurite outgrowth and neural differentiation compared to stiffer alternatives.[19] Encapsulation of human embryonic stem cell-derived neural stem cells (hESC-NSCs) within HA hydrogels improves cell viability and differentiation after spinal cord injury (SCI) while reducing cavity formation and glial scarring.[19] Additionally, HA-based electrospun scaffolds, such as HA/heparin nanofibers, accumulate growth factors like bFGF and NGF, promoting neuronal differentiation and enhancing cell proliferation.[20]

Gelatin is a denatured protein derived from collagen through acidic (type A) or alkaline (type B) hydrolysis.[21] Its properties vary depending on the collagen source, animal age, and processing methods.[22, 23] Gelatin provides a biocompatible environment that supports cell adhesion, growth, and differentiation, making it widely used in tissue engineering.[24–27] It is water-soluble and undergoes reversible gelation in response to temperature changes.[28] However, gelatin scaffolds can lose their structure in aqueous environments, necessitating modifications to improve their mechanical and thermal stability.[28, 29]

Gelatin-based hydrogels have shown great promise in neural tissue engineering, supporting the growth and viability of neural cells in 3D cultures.[19] Studies have demonstrated that gelatin scaffolds offer appropriate stiffness and stability, promoting cell proliferation and making them promising candidates for CNS repair and regeneration.[30]

Hydrogels have become materials capable of mimicking the tissue microenvironment, enabling controlled growth factor delivery, and being patternable for fabrication into desired constructs.[31] In the context of spinal cord and PNI, gelatin methacrylate (GelMA) has shown significant promise.[32] Recent studies have demonstrated using GelMA-based hydrogels in digital light processing (DLP) 3D printing to fabricate multichannel NGCs for peripheral nerve regeneration.[30] These GelMA-based NGCs provided an excellent environment for neural PC12 cells, supporting their survival, proliferation, and directional migration along the longitudinal channels. Furthermore, neural crest stem cells (NCSCs) cultured on the printed GelMA-NGCs were able to differentiate into neurons, highlighting the potential of GelMA-based NGCs in neural tissue engineering.

F11 DRG neural cells, derived from neonatal rat dorsal root ganglia (DRG) neurons fused with the mouse neuroblastoma cell line N18Tg2, serve as an established *in vitro* model for replicating features of injured peripheral neurons.[33] These cells exhibit peripheral sensory neuron characteristics, including a wide array of ion channels and specific neuronal markers, and are widely used to study signalling and neurite outgrowth. Their responsiveness to electromagnetic stimulation makes them ideal for exploring regenerative mechanisms. F11 cells have been employed in studies of neuronal plasticity and transcriptional regulation during PNS repair.

In material-based approaches, F11 cells have been exposed to pulsed electromagnetic fields (PEMFs) produced by a custom-made device.[34] By modulating parameters such as frequency, pulse repetition frequency, and duty cycle, optimal conditions for enhancing cell viability and neurite outgrowth were identified.[34] PEMFs have been shown to promote neural regeneration in F11 cells by accelerating neurogenesis, as confirmed by gene expression analysis 10 days post-stimulation.[34] Recent studies also demonstrate the protective effects of low-frequency, low-energy PEMFs on neuron-like and microglial cells under inflammation- and hypoxia-induced injury conditions.[35] In neuron-like cells (e.g., SH-SY5Y and PC12), PEMF exposure reduced hypoxia-induced damage by lowering cell death and oxidative stress. In microglial N9 cells, PEMF treatment decreased the release of pro-inflammatory cytokines (TNF-α, IL-1β, IL-6, IL-8) after lipopolysaccharide (LPS) stimulation, suggesting both neuroprotective and anti-inflammatory effects. These findings position PEMFs as a promising strategy for promoting neural regeneration and mitigating ischemic injury, potentially offering new translational approaches combining grafting with external field control in patients.

GelMA synthesized from type A gelatin has been shown to have no immunogenic effects on surrounding neural tissues *in vitro* and *ex vivo*.[32] While hyaluronic acid (HA) is widely used in neural tissue engineering, its application in neural conduits is relatively unexplored. In this study, we report the first combination of HAMA-GelMA hydrogels with growth factors to enhance the migration of neural cells. Additionally, the significant limitations of HAMA and GelMA, such as poor mechanical properties, were overcome by incorporating a unique 3D-printed polycaprolactone (PCL) fiber grid architecture. Hence, in this study, we fabricated a GrooveNeuroTube, incorporating 3D-printed PCL fibers and a composite HAMA-GelMA hydrogel loaded with bioactive agents (neural growth factor - NGF, glial cell line-derived neurotrophic factor - GDNF, vascular endothelial growth factor – VEGF, and lysozyme). The surface of PCL fibers was engineered with a specific micro-topography to guide neural cells and maintain conduit integrity. We hypothesized that the developed conduit, consisting of overlapping PCL fibers and an interconnected HAMA-GelMA hydrogel network, can significantly enhance the growth and migration of 3D bioprinted neural DRG cells, offering a promising strategy for nerve regeneration. For that, we utilized 3D bioprinting technology to mimic neural stumps, which require the placement of a neural conduit to enhance neural cell migration for regeneration. The cell-laden hydrogel, 3D bioprinted on both sides of the neural conduit, was cultured under prolonged conditions for 1 to 60 days to investigate the migration dynamics within the developed ***GrooveNeuroTube***. Furthermore, we specifically targeted the design of this conduit to support DRG F11 neuronal cell migration and neurite extension over prolonged periods, providing a new platform for ex vivo-like studies. Exemplary, we investigated in this model the synergistic effect of PEMF stimulation on neural regeneration, hypothesizing that the combined approach of bioengineered graft would significantly enhance cellular alignment, proliferation, and axonal growth.

## 2. Materials and Methods

### 2.1. PCL preparation

Polycaprolactone (PCL, Sigma Aldrich) pellet with a molar mass of M_n_ = 80 kDa was used for 3D printing.

### 2.2. 3D printing

#### 3D printing

Cellink Bio X2 - 3D printer (CellInk, Gothenburg, Sweden) was used for the PCL 3D printing. 33 mm (length) x 16.5 mm (width) rectangular constructs consisting of 2 layers of fibers were printed by pneumatic extrusion print head at a temperature of 90 °C. The optimization phase consisted of testing various printing temperatures (90 °C to 220 °C), different pressures (175–550 kPa), and printing speeds (10–15 mm/s), after which the following final printing parameters were selected for biological studies: 90°C, 180 kPa, and two printing speeds: 10 mm/s and 15 mm/s. For the 3D printing of the PCL, we used an aluminum cartridge and a metal nozzle with a pressure of 180 kPa to print directly onto a glass petri dish. After 3D printing, constructs were left to cool down at room temperature before being carefully removed. A 3D-printed PCL scaffold with dimensions of 33 mm (length) × 16.5 mm (width) was tightly rolled three times along its shorter side to form a tube with dimensions of 16.5 mm x approximately 7.5 mm (see ***Figure 1*** in the main text for more details). To secure the tube and prevent it from unrolling, a connecting section of the sheet was manually bonded to the tube by applying additional PCL fibers using the thermoplastic printhead of the BioX2, heated to 60°C.

**Figure 1:**
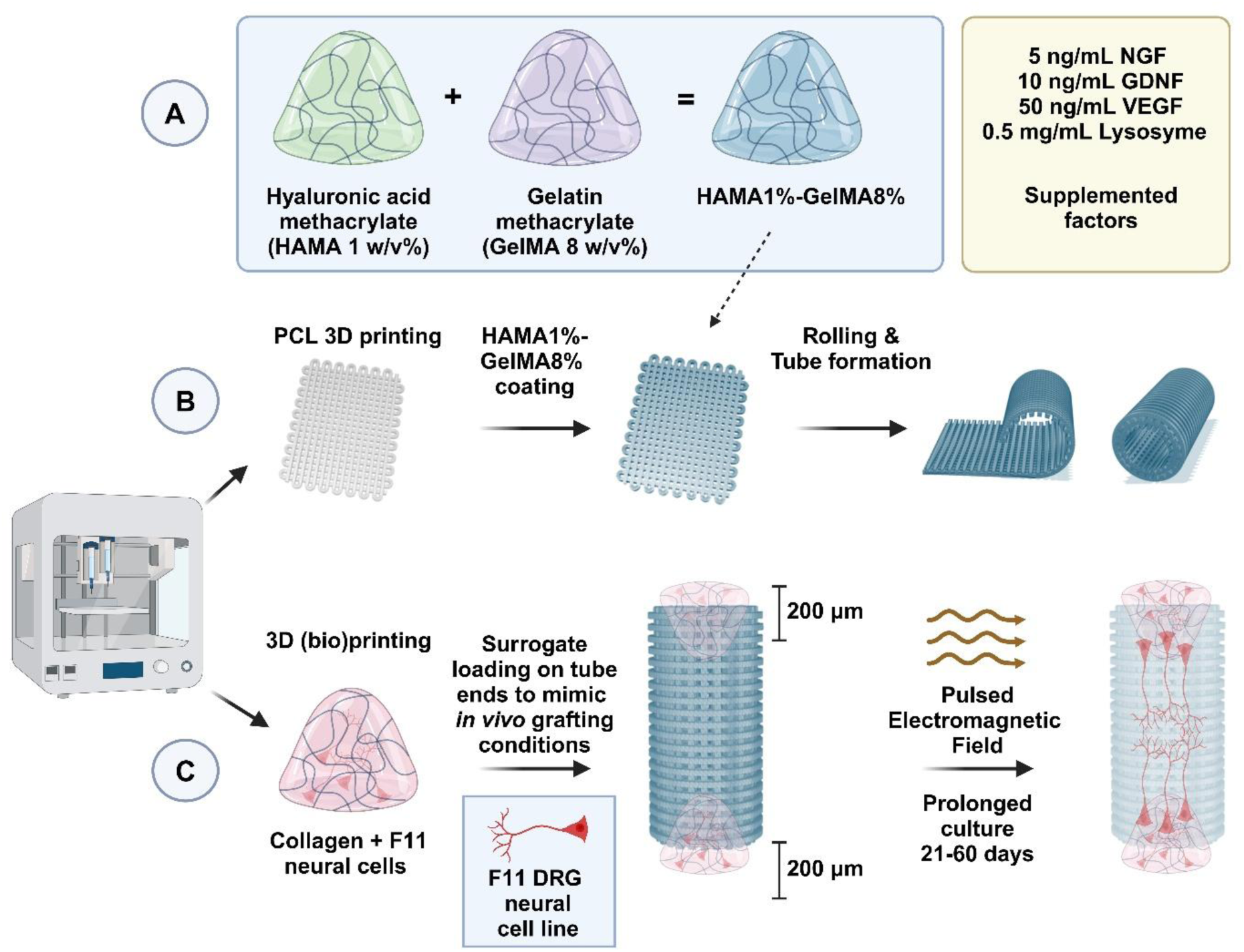
A) Preparation of the hydrogel formulation: Hyaluronic acid methacrylate (HAMA, 1 w/v%) is combined with gelatin methacrylate (GelMA, 8 w/v%) to produce the composite hydrogel, referred to as HAMA1%-GelMA8%. Selection of supplemented growth factors and antimicrobial lysozyme is noted in a separate table. B) Fabrication of the scaffold tube: A polycaprolactone (PCL) grid is first 3D-printed, followed by coating with the HAMA1%-GelMA8% hydrogel. Subsequently, the coated grid is rolled and formed into a tubular structure. C) 3D (Bio)printing and stimulation: Collagen mixed with F11 DRG neural cells are bioprinted at both ends of the tubular construct to mimic in vivo grafting conditions, spanning roughly 200 µm injection side into the tube. The structure can then be subjected to pulsed electromagnetic field (PEMF) stimulation and cultured for a prolonged period (21–60 days) to promote neural tissue outgrowth and integration. Created with BioRender.com.

### 2.3. Composite Hydrogel preparation

Methacrylated Gelatin (GelMA) powder with a high degree of methacrylation (DoM = 95%, PhotoGel® Catalog #5273, Advanced BioMatrix, Inc.) was dissolved in PBS preheated to 60°C and placed in a water bath with shaking for approximately 40 minutes until fully dissolved. For different experiments, hydrogels were prepared by dissolving GelMA powder in 1X PBS at 5, 10 or 15% (w/v) at 60°C with 37.4 mg/mL ruthenium photoinitiator and 119 mg/mL of sodium persulfate. Hydrogels were sterilized through a 0.2 µm syringe filter (83.1826.102, Starstedt). Hydrogels were photopolymerized using a 405 nm diode source (from Cellink BioX2) for 45 s (with power irradiation determined as 5.3 W at a 1 cm distance from hydrogel). Methacrylated Hyaluronic Acid (DoM = 45-65%, HAMA, PhotoHA®-Stiff Catalog #5275, Advanced BioMatrix, Inc.) powder with was added to cooled PBS (4°C) and left overnight in a refrigerator on a rotator to ensure complete dissolution. Once the GelMA solution was fully dissolved, it was cooled to 24°C. Meanwhile, the HAMA solution was warmed to 24°C and then combined with the GelMA solution using a positive displacement pipette. The mixture was placed on a rotor in a CO₂ incubator at 37°C for 30 minutes to form a uniform HAMA-GelMA suspension with 37.4 mg/mL ruthenium photoinitiator and 119 mg/mL of sodium persulfate. The concentrations of GelMA and HAMA were chosen such that the final concentrations of GelMA were 4, 8, or 10 w/v%, whilst the HAMA concentration was fixed at 1 w/v%. The preparation of the key hydrogel formulation used throughout manuscript is presented at ***Figure 1A***: Hyaluronic acid methacrylate (HAMA, 1 w/v%) is combined with gelatin methacrylate (GelMA, 8 w/v%) to produce the composite hydrogel, referred to as HAMA1%-GelMA8%. For other concentrations, the number in GelMA is simply updated in sample naming. In some cases, as described in the main text, the following bioactive compounds were added to the HAMA1%-GelMA8% composite at the mixing stage on a rotor in a CO₂ incubator at 37°C for 30 minutes: Nerve growth factor (NGF) at 5 ng/mL, Glial cell line-derived neurotrophic factor (GDNF) at 10 ng/mL, and Vascular endothelial growth factor (VEGF) at 50 ng/mL, along with lysozyme at 0.5 mg/mL.

### 2.4. Production of porous GroveNeuroTube - HAMA-GelMA 3D PCL tubes

To fabricate the composite HAMA-GelMA-PCL, a 3D-printed PCL grid with dimensions of 33 mm x 16.5 mm was laid flat in a Petri dish. Next, a preheated (to 24°C) solution of HAMA1%-GelMA8% was injected into the center of the PCL grid, allowing the solution to evenly distribute across the grid. At this stage, photopolymerization was initiated by exposing the hydrogel to a 405 nm diode light source (Cellink BioX2) for 45 seconds, with a power irradiation of 5.3 W at a 1 cm distance. Subsequently, the composite hydrogel containing the PCL was tightly rolled three times to form a tube measuring 16.5 mm in length and 5 mm in diameter, similarly to the protocol described above for PCL alone. To secure the structure and prevent unrolling, a connecting section of the sheet was again bonded to the tube by applying PCL fibers using the thermoplastic printhead of the BioX2, heated to 60°C. Once completely rolled, the tube was allowed to air-dry for 1 hour (***Figure 1B, C***). HAMA1%-GelMA8% containing growth factors (abbreviated as HAMA1%-GelMA8%-GF) was only added once the tube cooled down to 37°C.

### 2.5. Scanning electron microscopy (SEM)

The morphology of all samples was analysed using an SEM-7001TTLS microscope (JEOL Ltd., Akishima, Japan) operated at an accelerating voltage of 15 kV. Hydrogels with 5, 8, and 10% GelMA concentrations were further examined using the same SEM coupled with a PP3000T cryo-preparation system (Quorum Technologies, Laughton, UK). This setup enabled the preparation, processing, and transfer of cryo-fixed specimens directly into the SEM chamber for observation in a vitrified state. The samples were mounted on metal holders and rapidly cryo-fixed by immersion in sub-cooled liquid nitrogen (N_2_ slush) at −210 °C. The frozen samples were then transferred under vacuum into the SEM preparation chamber, maintained at −180 °C. Within this chamber, the specimens were fractured to reveal fresh inner surfaces, sublimated to remove surface ice, and coated with a thin layer of platinum. Subsequently, the samples were transferred under vacuum to the SEM cryo-stage, maintained at −190 °C, where imaging of the surfaces was performed. Additionally, elemental analysis of the cryo-prepared hydrogels was conducted at an accelerating voltage of 15 kV. Pore size analysis was performed on over 400 pores using ImageJ.

### 2.6. Fourier transform infrared spectroscopy (FTIR)

FTIR measurements were performed at room temperature between 400-4000 cm^−1^ using a Nicolet iS50 FT-IT spectrophotometer (Thermo Scientific, Waltham, United States). The HAMA1%, GelMA8%, HAMA1%-GelMA8%, and crosslinked HAMA1%-GelMA8% freeze-dried samples were placed onto the ATR crystal and pressed down to ensure contact between the sample and the crystal. The measurements were taken with a resolution of 4 cm^-1^ and for 64 scans.

### 2.7. Nuclear Magnetic Resonance (NMR)

To study the crosslinking process, ^1^H NMR spectra were recorded using an Agilent DD2 800 spectrometer (Agilent Technologies, Santa Clara, CA, USA) equipped with a 5 mm 1H/13C/15N probe. Measurements were conducted at 37 °C with D₂O as the solvent. The sample concentration was maintained at 80 mg/mL of GelMA (8%), 10 mg/mL of HAMA 1%, 0.18 mg/mL of ruthenium photoinitiator, and 0.56 mg/mL of sodium persulfate. The sample was exposed to UV light directly in an NMR tube to induce crosslinking. ^1^H NMR experiments were performed using Agilent’s standard pulse sequence with a low-power pre-saturation pulse for residual water suppression. To enhance the signal-to-noise ratio, 128 scans were accumulated per spectrum. The spectra were referenced relative to 3-(Trimethylsilyl)propionic acid-d₄ sodium salt (TSP) as an external standard.

### 2.9 Degradation and swelling

To evaluate the in vitro degradation behavior of the hybrid 3D conduit composed of 3D-printed PCL mesh and photocrosslinked hydrogel (GelMA 8% w/v and HAMA 1% w/v), we conducted a longitudinal mass loss analysis under simulated physiological conditions. Cylindrical constructs (length 16.5 mm, outer diameter 7.5 mm) were fully immersed in phosphate-buffered saline (PBS) and incubated at 37°C in a 5% CO₂ atmosphere. The constructs (n = 3) were first weighed (analytical balance) at day 0 and subsequently re-weighed at 3, 7, 14, 28, and 60 days following removal, PBS removal, and drying (blotting + short desiccation).

The swelling capacity (S%) was determined using gravimetric method. Initially, the dry weigh of each sample (W_o_) was measured followed by immersion of the samples in 10 mL of Milli-Q water and retrieved at predetermined intervals upto 720 min. After removal, excess surface water was gently blotted with moistened filter paper and the samples were weighed again (W_f_). The swelling capacity was calculated using equation (1)

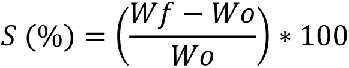

### 2.9. Rheological measurements

The storage (G’) and loss (G”) moduli of the hydrogels were measured using an TA Instruments AR 2000ex rheometer equipped with a 12 mm diameter top plate and a 1 mm gap in a parallel plate geometry. For sample preparation, 500 μL of hydrogel was photopolymerized in a specially designed ceramic mould with wells measuring 20 mm in diameter and 10 mm in height, sealed at the bottom with a removable PDMS layer. The samples were carefully transferred onto the stationary bottom plate of the rheometer using a spatula, ensuring minimal disturbance to the sample structure. The top plate was gradually lowered to avoid disrupting the hydrogel. Before each measurement, the sample was equilibrated at 37 °C for 3 minutes. Strain sweeps were performed from 1% to 10% strain at a fixed frequency of 1 Hz, and frequency sweeps were conducted from 1 Hz to 20 Hz at a constant strain of 1% within the linear viscoelastic region. Measurements included G’, G’’ and complex viscosity |η*|. Each measurement was repeated three times to ensure reproducibility.

### 2.10. Cell Culture

Immortalized Dorsal Root Ganglion Neuron Cell Line (F11 cell line from rat, 08062601, ECACC) were purchased from Merck and cultured in DMEM + 2mM Glutamine + 10% Foetal Bovine Serum (FBS). The medium was supplemented with penicillin (100 U/mL) and streptomycin (100 mg/mL), and F11 were cultured at 37°C in a humidified atmosphere of 5% CO_2_. Cells were grown in sterile 25 cm^2^ and 75 cm^2^ tissue culture flasks. When cells reached 80% confluence, they were transferred and sub-cultured in a new flask by trypsinization with 0.05% trypsin. Cell growth and confluence were observed under the light microscope and counted by an Automated Cell Counter (BioRad). Experiments were carried out with cells from the third to sixteen passages.

### 2.11. DRG neuronal cells seeding prior to microscopies imaging

PCL fibers pattern matrices were sterilized by 70% ethanol washing followed by subsequent 30 min UV light exposure (λ=254 nm). Samples were exposed to 1 hour of UV light exposure and placed in sterile multi-well plates. After reaching 80% confluency, the F11 cells were washed with Dulbecco’s Phosphate Buffered Saline containing no calcium, magnesium, or phenol red (D-PBS, ThermoFisher, 12604-013) and collected by trypsinization at 37 °C for 5 minutes. Cells were counted by Automated Cell Counter (BioRad). Dissociated F11 cells were plated onto PCL grids at a density of 25 × 10^4^ cells per cm^2^. The medium was replaced at 3 days post-seeding and then refreshed every 3–4 days. Cells were grown in humidified conditions (95% humidity) at 37°C and 5% CO_2_.

### 2.12. 3D bioprinting on GrooveNeuroTube – *ex vivo* model

10X PBS or 10X culture medium chilled to 15 °C was combined with a 8 mg/mL methacrylated type I bovine collagen solution (PhotoCol®, Catalog #5198, Advanced BioMatrix, Inc.)) chilled to 4 °C in an 8:1 ratio under gentle swirling. The pH was carefully adjusted to 7.0–7.5 using sterile 0.1 M NaOH and monitored with a pH meter or phenol red. The volume was brought to 10 parts with sterile water, ensuring the mixture’s temperature remained between 2–10°C to prevent premature gelation. The mixture was then warmed to 37°C and allowed to form a gel over 90–120 minutes. For gels containing F11 DRG neural cells, a 1:1 (v/v) mixture of a cell suspension (500,000 cells/mL) and collagen was prepared at 4 °C. Aliquots (0.5 mL) were placed into a 24-well plate, with one column left as a non-crosslinked control, and microbial transglutaminase (TG) (3.33:1 collagen: TG w/w%) added to the others. Plates were prepared for each time point and incubated under standard culture conditions, and media changes were made every 3 days. Time-zero samples were analyzed immediately after a 30-minute incubation. The pre-printed tube was used as a base for bioprinting collagen. The bioink, consisting of F11 DRG neural cells pellet embedded in collagen, was loaded into the bioprinter cartridge. Subsequently, 1 layer of this bio-ink of 500 μL of cell-laden (250,000 cells) collagen was deposited onto one end of the tube within 3 minutes at room temperature using a 20G cylindrical needle and pressure of 15 kPa. Afterward, bioprinting of the equivalent volume of the bio-ink was performed on the opposite end of the tube. The tubes were then placed horizontally in 6-well plates and incubated to allow mTG crosslinking. After 30 minutes, medium was added to the wells, and a medium change was performed the following day.

### 2.13. Migration test – 3D cell bioprinting on both tube ends

Migration within 3D tubes was evaluated using confocal microscopy and cell labeling. To estimate cell viability and migration within the constructs, the Fluorescein Diacetate/Propidium Iodide (FDA/PI) viability assay was employed. Constructs were sectioned into 500 µm transverse slices using a vibratome at three time points: day 1 (at the gel corner), day 14, and day 28 of culturing (from the central region) (as shown in main text in ***Figure 8***). Samples were embedded in 6% agarose blocks (Condalab, 8017.00) and sectioned transversely at 500 µm thickness using a vibratome (Leica VT100S). The sections were immersed in DMEM medium without foetal calf serum (FCS), containing FDA (5 mg/mL) and PI (2 mg/mL), for 5 minutes. Following incubation, the staining solution was removed, and the cross-sections were washed twice with PBS. The samples were then placed in DMEM medium without FCS and immediately observed using a IX 73 Olympus for live/dead imaging. The remaining constructs were used for immunostaining. Constructs were fixed in 4% paraformaldehyde overnight at 4°C. The next day, tissues were washed three times for 30 minutes each with 1X PBS on a rotary shaker. Samples were embedded in 6% agarose blocks (Condalab, 8017.00) and sectioned transversely at 300 µm thickness using a vibratome (Leica VT100S). Sections were incubated for 4 hours in blocking buffer (1X PBS, 0.5% Triton X-100, 1% BSA). Primary antibodies were used to detect antigens: rabbit anti-Dcx (1:100, ThermoFisher), mouse anti-Map2 (1:250, ThermoFisher), and rabbit anti-Syp (1:100, Abcam). After PBS washing, secondary antibodies such as Donkey anti-mouse Alexa Fluor 488, Donkey anti-rabbit Alexa Fluor 555, and others were incubated for 1 hour at room temperature. DAPI (1:10000, Molecular Probes) was used for nuclear staining. Based on images, cell orientation was analyzed using the “OrientationJ” plugin in Fiji. Normalization was done to the sum of all values. The orientation charts were created using Python’s “Matplotlib” library to visualize data in a polar coordinate system. The data, consisting of angles and corresponding values, was loaded from a file based on the measurements from Fiji. To enhance visual clarity, a polar bar plot was generated with customized colors, borders, and transparency. The plot’s axis was adjusted to display angles from -90° to 90°, with the zero-angle positioned at the top.

To estimate cell migration and distribution throughout the conduit, we performed fluorescent cell labeling. Cells used for seeding at one end of the conduit were labeled with CellTrace™ Far Red DDAO-SE (C34564, Thermo Fisher), while cells seeded at the opposite end were labeled with the CellTrace™ CFSE Cell Proliferation Kit (C34554, Thermo Fisher). Specifically, F11 cells were stained with 10 μM of CellTrace™ Green CMFDA and CellTracker™ Deep Red according to the manufacturer’s protocol. Briefly, cells were incubated with the fluorescent dyes (1:1000 dilution in DMEM) for 20 minutes at 37 °C, followed by five washes with PBS. After staining, cells were centrifuged at 400 × g for 5 minutes, resuspended in 10 mL of complete medium, and centrifuged again. The labeled cells were then used to prepare the collagen-mTG bioink, as described in section 2.11. Each stained cell population was used to bioprint into opposite ends of the GrooveNeuroTube. Specifically, 500 μL of collagen-based bioink containing 250,000 labelled-cells was bioprinted onto one end of the conduit as described in 2.11. The same procedure was repeated for the opposite end using the second cell population. After 28 days of culture under standard and PEMF conditions, constructs were fixed in 4% paraformaldehyde overnight at 4 °C. The next day, the samples were washed three times (30 minutes each) in PBS on a rotary shaker, embedded in 6% agarose (Condalab, 8017.00), and sectioned transversely into 300 μm slices using a vibratome (Leica VT100S). The central sections were immersed in PBS and stained with DAPI (1:10,000; Molecular Probes) for nuclear visualization. Fluorescence imaging was performed using a confocal microscope (Olympus FV1200) with a 10× dry objective and excitation lasers at 405 nm, 488 nm, and 635 nm. Emission signals were collected in the ranges of 430–460 nm for DAPI (excited at 405 nm), 510–520 nm for CFSE (excited at 488 nm), and 630–640 nm for CellTrace™ Far Red (excited at 635 nm). A 10×/NA 1.2 dry objective was used for imaging. After 28 days in culture, fluorescence imaging clearly showed co-localization of both labeled cell populations in the central region of the conduit, confirming that cells seeded from opposite ends successfully migrated and met in the middle. All experiments were performed with four biological repeats and three experimental repeats.

### 2.14. Cytotoxicity assessment

For live/dead visualization, sections were incubated with 2 μM Calcein-AM and 2 μM Ethidium homodimer-1 in PBS at 37°C for 30 minutes. The cells were visualized on confocal laser scanning microscope (Zeiss LSM 780 NLO, Olympus FV1200) with a 40× water objective and excitation laser wavelengths of 561 nm for EthD-1 and 488 nm for Calcein-AM and emission of 500-520 nm and 600–630 nm, respectively. The live-to-dead cell ratio from live/dead measurements was calculated by dividing the number of live cells by the total cell count (sum of live and dead cells) and expressing it as a percentage for each time point. The percentage of dead cells was obtained by subtracting the percentage of live cells from 100%. Next, the increase in live cell growth was calculated relative to day 1 by normalizing the live cell count at each subsequent time point to the initial value. The number of live cells on day 1 was set as 100%, and the percentage increase was calculated using the formula: Live Cell Growth (%) = (Live Cells at given time point / Live Cells at day 1) × 100%. These calculations were performed to assess cell viability and proliferation trends over time, including the effects of PEMF stimulation. To perform a quantitative viability assay using flow cytometry (MUSE), the commercial Muse™ Annexin V & Dead Cell Kit was used. For sample preparation, the cell cultures after 14 days of incubation with respective materials were washed with PBS, trypsinized, centrifuged, and resuspended in 100 µL of PBS with 1% FBS. Then, 100 µL of the reagent was added, and the mixture was incubated for 20 minutes in the dark at room temperature. Finally, the collected data were analysed using Flowing Software 2. Another experiment conducted was the detection of reactive oxygen species (ROS) using the Muse® Oxidative Stress Kit. Cells were collected from the culture in the same manner as for the viability test. The positive control consisted of cells seeded the day before and treated with menadione one hour prior to measurement. After the measurements, the results were analysed using Flowing Software 2.

### 2.15. Immunocytochemistry and quantification

Cells cultured on all surfaces for 2, 7, 14, and 21 days were rinsed twice with PBS (phosphate-buffered saline, Sigma-Aldrich) and incubated for 15 min with 2.5% formaldehyde solution in PBS at T = 21 °C. Then, the cells were permeabilized by PBS containing 0.5% Triton X-100, washed with PBS, blocked with 3% BSA in PBS, and incubated with primary antibodies for 1 hour at room temperature. The following primary antibodies were used to detect antigens: rabbit anti-Dcx (IgG, 1:100, 48-1200, ThermoFisher), mouse anti-Map2 (IgG1, 1:250, 13-1500, ThermoFisher), rabbit anti-Syp (IgG, 1:100, ab32127, Abcam). After washing with PBS, the secondary antibodies were used to react with the cells for 1 hour at room temperature. The following secondary antibodies were used: Donkey anti-mouse (Alexa Fluor 488, 1:200, a24350, ThermoFisher), Goat anti-rabbit IgG (H+L) (Alexa Fluor 647, a32733, ThermoFisher), Goat anti-mouse IgG (Alexa Fluor 488, a32723, ThermoFisher). Finally, cells were stained by 4’,6-diamidyno-2-fenyloindol (DAPI) and imaged immediately using Zeiss LSM 780 NLO and Olympus FV1200 confocal microscope with 40× microscope water objective. All measurements were repeated at least three times. The MAP2/SYP/DCX positive cells were determined manually from at least 500 cells taken from random immunocytochemical (ICC) staining. DRG cells were quantified by counting the number of MAP2+ and/or SYP/DCX+ cells possessing a neuronal morphology expressed as a percentage of DAPI-stained nuclei per condition.[36] Neurite outgrowth was characterized by measuring the most extended neurite length for each MAP2/SYP positive cell. As reported previously, at least 100 DCX/MAP2/SYP positive cells per sample were included in the measurement.[37]

### 2.16. RNA isolation and RT-qPCR

The series of real-time polymerase chain reaction (qPCR) experiments were conducted on F11 DRG neural cells using designed primers (***Table S1***). Obtained cell suspensions were used for RNA isolation. RNA was isolated using the Aurum Total RNA Mini Kit (Bio-Rad), yielding >500 ng/μL of RNA. During the isolation, DNase treatment was performed to eliminate any remaining genomic DNA that could potentially amplify during PCR reactions. Subsequently, reverse transcription reactions were performed using 1 μg of RNA and the iScript cDNA Synthesis Kit (Bio-Rad) in a volume of 20 μL. The cDNA synthesis reaction conditions were as follows: 5 min at 25°C, 20 min at 46°C, 1 min at 95°C. The obtained cDNA served as a template for PCR reactions using gene-specific primers, including GAPDH, ACTB, DCX, MAP2, TUBB3, SYP, SYN, s100b, and peripherin. The cDNA samples were diluted 2.5x and reaction was carried out using 2 μL of cDNA template and Taq PCR Master Mix (Qiagen). The exemplary PCR reaction conditions were as follows: initial denaturation at 94°C for 3 min, denaturation at 94°C for 30 s, primer annealing at 60°C for 30 s, extension at 72°C for 60 s (35 cycles), and final extension at 72°C for 10 min. The PCR products were separated on a 1.5% agarose gel, and by comparing them to a size marker (Puc19/msp1), specific products corresponding to 104 bp for ACTB, 158 bp for GAPDH, 151 bp for MAP2, and 89 bp for TUBB3 were confirmed. The PCR products for ACTB, GAPDH, MAP2, and TUBB3 were excised from the gel and purified using the QIAquick Gel Extraction Kit (Qiagen). The amounts of DNA obtained for each sample were quantified in ng/μL and are presented in Supplementary materials. Subsequently, molecular quantities per μL were calculated based on the product size and DNA concentration, and decimal dilutions were performed.

### 2.17. Pulsed electromagnetic fields (PEMF) device and stimulation

For the PEMF tests, we used an in-house stimulation system (***Figure S7***). The system could generate sinusoidal, triangle, and square waveform functions in the 0.1 Hz - 10 MHz frequency range. A smaller 50-500 Hz range was used as a sinusoidal wave for cell stimulation. Synthesized Function Generator SFG-2000/SFG-2100 Series (GW INSTEK) was used to generate alternating current: sinusoidal, triangle, and square-shaped of various frequencies. The shape of the curves was confirmed on the MDO-2000E series, which is a multi-functional mixed-domain oscilloscope (GW INSTEK). Curer fed the six ferrite core coils (Viston art. no. 3808) under the 6-well polystyrene transparent multiwall plate. The magnetic field obtained was measured on the bottom of the wells, and a magnetic field meter (TENMARS TM-197) was used for AC magnetic measurements. The sterile closed cell culture in Petri dishes was placed directly above the coils, providing a magnetic field. For the PEMF, we chose a sinusoidal function of frequency 50 and 500 Hz, which generated magnetic field induction of 0.24 mT at the bottom of the well. The device was used to stimulate cells 4 h daily in the incubator for the whole duration of the experiment.

### 2.18. Antibacterial tests

The optical density measurements at 600 nm (OD600) were carried out to monitor the growth of microbes, i.e. *E.coli* and *S.aureus* in a liquid culture. The overnight cultures of 150 colony-forming units (CFU)/mL *S.aureus* (150 μL) or *E.coli* (150 μL) were added to the tubes containing 30 mL of fresh Luria broth (LB) medium. Discs of crosslinked HAMA1%-GelMA8% containing lysozyme (0.5 mg/mL) (5 mm x 5 mm) were incubated for 1, 7, 14 and 21 days. Bacterial strains *E. coli* ATCC 35218 and *S. aureus* ATCC 29213 were grown in LB Broth Lennox under continuous shaking at 250 rpm, at 37 °C, for 24 hours. Following incubation, the bacterial cultures were diluted to achieve a final 5 × 10^5^ cells/mL concentration in LB broth. A volume of 100 μL of the prepared bacterial suspension was transferred into each well of a 96-well plate. Fresh solutions of lysozyme at concentrations of 0.25, 0.5, 1, and 1.5 mg/mL were prepared and incubated for 1, 7, 14, and 21 days to simulate conditions resembling those found in hydrogels stored/incubated in the incubator. These pre-incubated lysozyme solutions were introduced into the wells alongside the bacterial cultures and incubated at 37 °C for 24 hours. The bacterial growth was assessed by measuring the optical density (OD600) using an Infinite M200 Pro microplate reader (Tecan) after incubation. To minimize any potential interference caused by the light-scattering properties of the samples, control wells containing the same concentrations of lysozyme in LB broth but without bacteria were included as blank controls. For comparison, untreated bacterial cultures served as the negative control, while bacterial cells exposed to antibiotics were used as the positive control. All experiments were conducted in triplicate to ensure reproducibility and reliability of the data.

Subsequently, a 0.5 mg/mL lysozyme concentration was selected for further studies. The HAMA-GelMA hydrogel containing 0.5 mg/mL lysozyme was prepared as described in the previous sections. The hydrogel, immersed in PBS, was placed in an incubator for the duration of the experiment. After 7, 14, and 21 days of incubation, 100 μL of the PBS supernatant was collected, and bacteria at the same concentration as in the previous experiment were added to the PBS supernatant containing lysozyme released from the HAMA-GelMA hydrogel. A volume of 100 μL of the prepared bacterial suspension mixed with the lysozyme-containing PBS supernatant was transferred into each well of a 96-well plate. Optical density (OD600) measurements were taken after 24 hours, following the same procedure as in the previous experiment.

## 3. Results and Discussion

Peripheral nerve injuries are challenging to treat due to the slow regeneration rate and limited functional recovery. Current NGCs often fail to replicate the biomimetic microenvironment required for effective nerve repair. In this study, we developed a tubular nerve conduit using a combination of 3D printing (***Figure 1B***) and bioengineered bioactive hydrogel (***Figure 1A-C***) as a new *ex-vivo* model. Firstly, we formulated a composite hydrogel consisting of hyaluronic acid methacrylate (HAMA, 1 w/v%) and gelatin methacrylate (GelMA, 8 w/v%), denoted as HAMA1%-GelMA8% (***Figure 1A***). A 3D-printed polycaprolactone (PCL) grid scaffold was coated with this hydrogel, rolled into a tubular structure, and used as a nerve conduit (***Figure 1B***). In a final step, cell-laden collagen was bio-printed onto both ends of the tube to mimic *in vivo* grafting conditions. Subsequently, to enhance neural regeneration, the conduit was subjected to a PEMF stimulation during prolonged culture periods (21–60 days) (***Figure 1.C***). The hypothesis was to test our developed *ex vivo*-like system and check if PEMF stimulation supported neural cell migration, alignment, and outgrowth, mimicking the *in vivo* conditions.

### 3.1 Fabrication of PCL fibers grid

In this study, following previous reports on PCL groove-patterned surface topography and its impact on neural cell differentiation and maturation [7, 38] we expand on this by exploring the influence of PCL fibers and the stiffness of HAMA-GelMA hydrogel on the growth and migration of neural DRG cells.

In the first part, we focussed on PCL fibers grid optimization. For this, we used various printing temperatures (90 °C to 220 °C), different pressures (175–550 kPa), and printing speeds (10–15 mm/s) to achieve consistent fibers with homogenous and reproducible layer-by-layer grid structure. After the preliminary elimination of challenges (***Figure S1***), we selected the following parameters for further studies: 90 °C, 180 kPa, and two printing speeds: 10 mm/s (abbreviated as 10N) and 15 mm/s (abbreviated as 15N). Both selected printing speeds enabled the generation of reproducible fibers (***Figure 2A***). For samples printed at 10 mm/s, the average fiber diameter was 96.1 ± 18.3 μm. At 15 mm/s, the average fiber diameter was 79.5 ± 19.7 μm (***Figure 2 A, E***), where the square size between all fibrillar constructs defines porosity. After printing two layers of PCL fiber grids, the structure was rolled into a tube (***Figure 2B, D***). The PCL fibers were tightly rolled three times to form a tube measuring 16.5 mm long and 7.4 mm in diameter. To secure the structure and prevent it from unrolling, a section of the sheet was bonded to the tube by applying additional PCL fibers using the thermoplastic printhead of the BioX2, heated to 60°C (***Figure 2 B, D***). We then investigated the topography of the produced PCL fibers and grids using optical microscopy and scanning electron microscopy (SEM). We confirmed it formed a regular structure (***Figure 2 A-D***).

**Figure 2:**
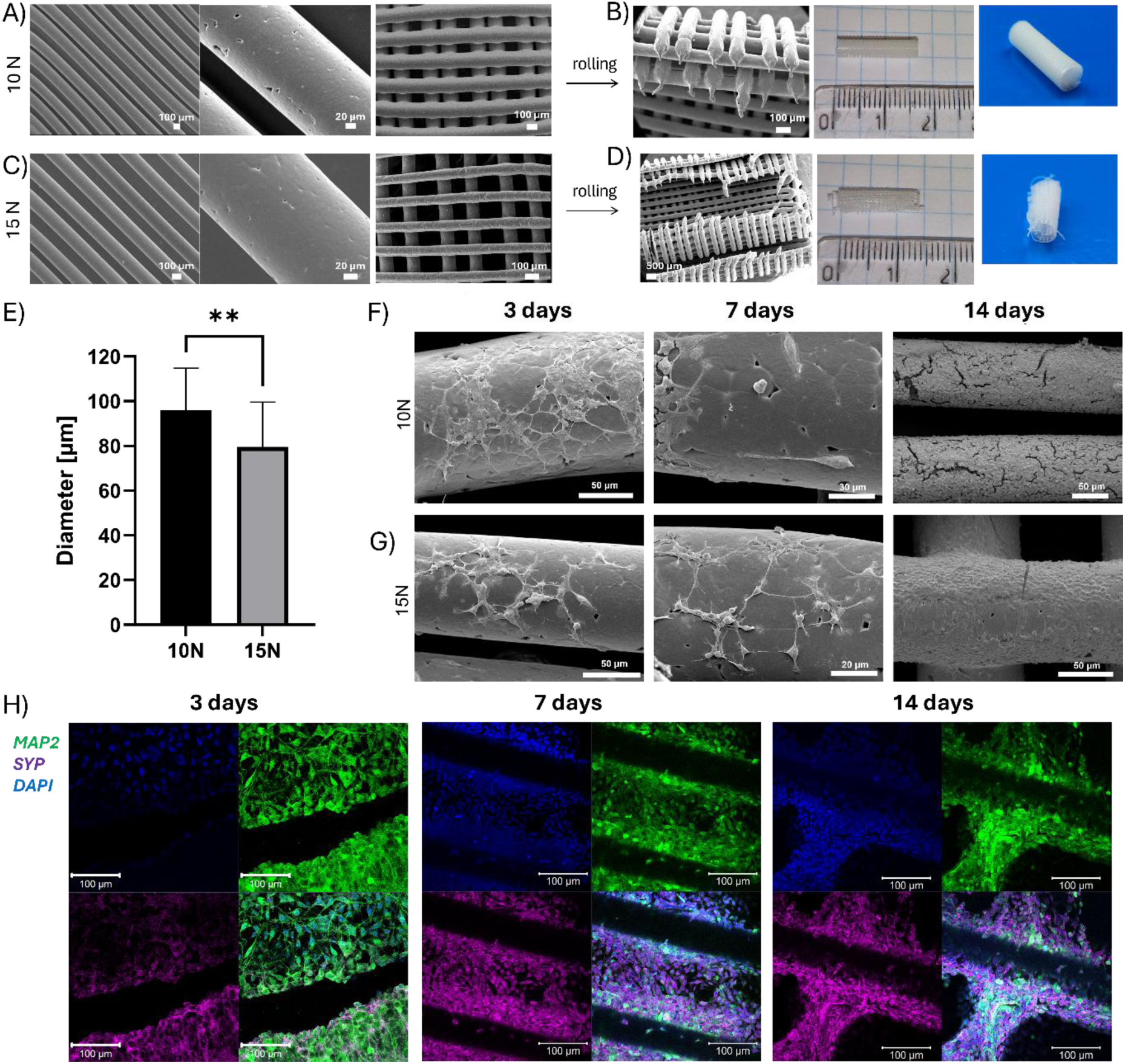
Optimization of 3D printing of PCL fibers: A) 10N and C) 15N, B, D) SEM morphological characterization of 3D printed PCL grid which we used to form multilayer tube. E) diameter of PCL fibers, F, G) SEM images of F11 DRG neural cells cultured on a PCL grid for 3,7, and 14 days. H) Confocal images of F11 DRG neural cells cultured on PCL showing the detection of late neuronal (green, MAP2) and synaptic vesicles (violet, SYP) markers.

Subsequently, we cultured the F11 DRG neural cell line on the fabricated PCL grids for 3, 7, and 14 days and assessed their morphology using SEM (***Figure 2F, G***). On day 3, we observed that F11 cells cultured on larger N10 fibers exhibited a noticeably higher cell density than smaller N15 fibers, likely due to the increased surface area of the N10 fibers. By day 7, F11 cells displayed a polarized, elongated morphology, indicating their alignment and adaptation to the scaffold structure. By day 14, the cells had formed extensive neurites and a dense cellular network on both types of fibers, with a visibly more robust network observed on the N10 grid, further emphasizing the advantages of the larger fiber diameter for neural cell growth and improved cell-cell connectivity (***Figure 2F, G***). From the two choices, the 10N PCL grid proved to be more stable, easier to roll and retained more coherent smooth structure (***Figure 2A-D***). It also appeared to provide better F11 DRG neural cells maturation (***Figure 2F***). Therefore, for all the subsequent evaluations we selected 10N PCL grids and tubes.

To confirm the functional process of neuronal maturation, we performed immunocytochemistry (ICC) to detect MAP2 (Microtubule-Associated Protein 2), a marker of mature neurons, and SYP (Synaptophysin), a protein characteristic of synaptic vesicles and considered a synaptic marker. MAP2+/SYP+ cells were identified across both grid types ***(Figure 2H***). With extended culture duration, the number of SYP+ cells increased, indicating enhanced synaptic activity, while MAP2+ levels remained relatively constant. The most prominently elongated MAP2+/SYP+ cells were observed on day 14, often in close contact with the neighbouring cells. This observation suggests the potential formation of neural-like networks, indicating ongoing neuronal maturation (***Figure 2H***).

### 3.2 Development of HAMA-GelMA composite hydrogel

Next, we evaluated the selection of composite hydrogels made from HAMA and GelMA. In similar composite formation, HAMA concentration was fixed at 1%, whereas secondary polymer concentration modulated to obtain desired properties, which we decided to follow through.[39] ***Figure 3A*** presents a schematic illustration of the composite hydrogel formation, showing the chemical composition of the two chosen polymers, HAMA and GelMA. In the first instance, we performed Cryo-SEM imaging of HAMA1%-GelMAX% hydrogels, where X indicates variable content of GelMA, going from 5, 8 to 10% (***Figure 3B-D***). All micrographs demonstrated interconnected porous structures of hydrogels. HAMA1%-GelMA8% displayed the most homogenous structure across all tested hydrogels, with the average porosity obtained from lognormal distributed in the 0.81 ± 1.82 µm range (***Figure 3E-G***). This was much smaller than for the HAMA1%-GelMA5%, for which a non-standard broader distribution that could not fit with any standard model was obtained.

**Figure 3:**
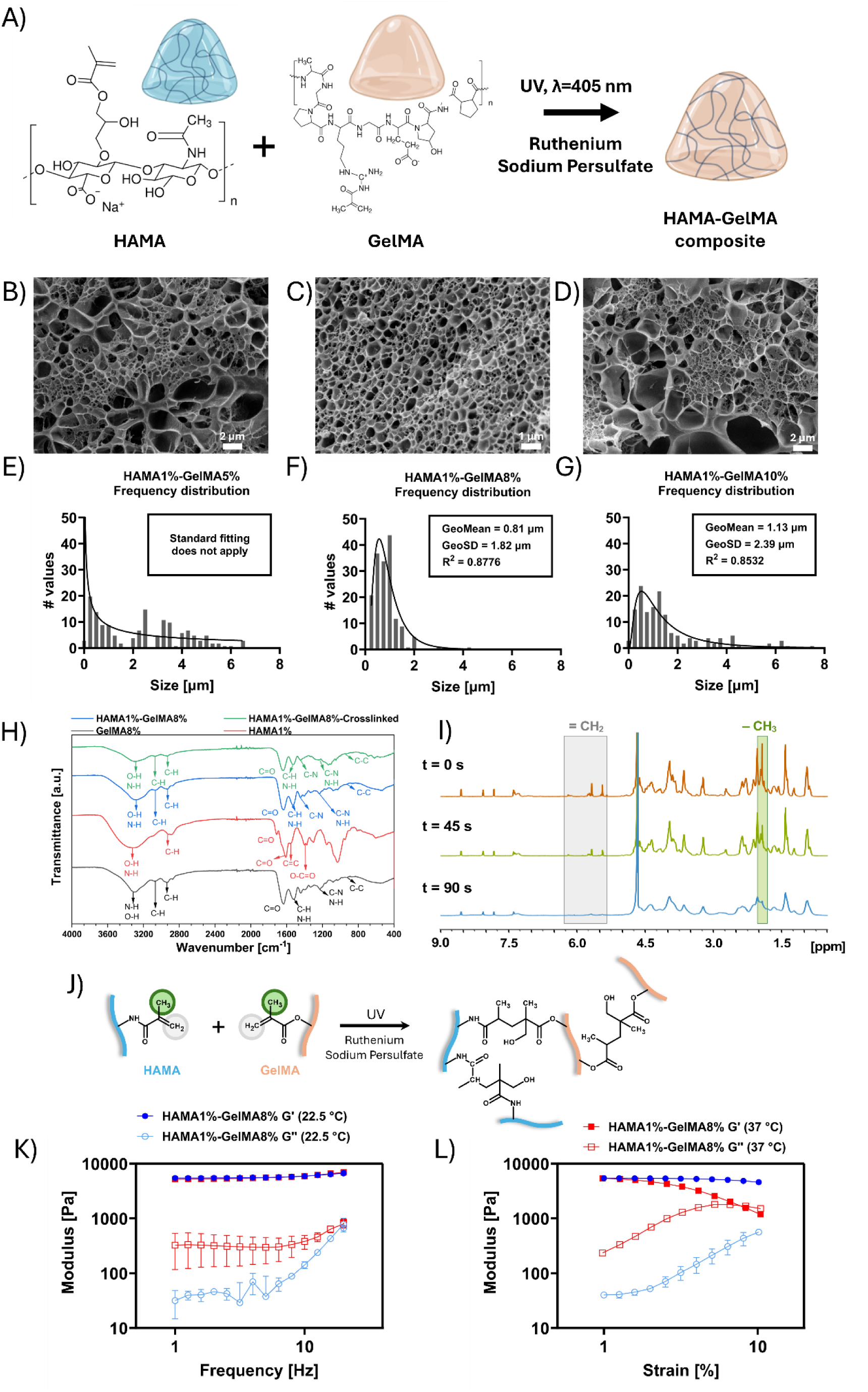
A) Scheme of HAMA-GelMA composite formation. CryoSEM of B) HAMA1%-GelMA 5% C) HAMA1%-GelMA 8% D) HAMA1%-GelMA10% E,F,G) pore size frequency distribution of each associated formulation H) FTIR spectra of HAMA, GelMA, and HAMA1%-GelMA8% composite hydrogels; I) ^1^H NMR spectra of non-crosslinked and crosslinked HAMA1%, GelMA8%, and HAMA1%-GelMA8% at different time points (note reduced content of crosslinker and initiator in methods to enable monitoring of the reaction); J) Schematic representation of the photoinitiated crosslinking reaction between HAMA and GelMA under UV irradiation in the presence of a redox initiator system; K) Amplitude sweep and L) frequency sweep of the HAMA1%-GelMA8% composite

The FTIR spectra of GelMA8% in (***Figure 3-H***) show several characteristic peaks. The broad peak 3300 cm^-1^ represents the stretching vibrations of O-H and N-H groups. Peaks within the range of 3100–2800 cm^-1^ are attributed to C-H stretching vibrations. The signature amide bonds of gelatin are evident at 1630 cm^-1^, corresponding to C=O stretching of amide I, 1520 cm^-1^ attributed to N-H bending coupled with C-H stretching of amide II, 1226 cm^-1^ signifies the C-N stretching and N-H bending of amide III, and 926 cm^-1^ indicating -C-C-skeletal stretch of amide IV.[40, 41]

In HAMA1% spectra broad peak around 3300 cm^-1^ indicate the stretching vibrations of O-H and N-H, while the peak at 2900 cm^-1^ represents C-H stretching. The region between 1710 cm^-1^ and 1450 cm^-1^ shows overlapping of amide I and amide II peaks along with C=O bending vibrations.[42–44] Peaks at 1553 cm^-1^ and 1290 cm^-1^ indicate stretching vibrations of C=C and O-C of methacrylate.[45] Additionally, the peak at 1410 cm^-1^ corresponds to stretching vibrations of O-C=O groups.[46]

In the composite HAMA1%-GelMA8% hydrogel, the dominant peaks of GelMA due to its higher concertation are observed along with the characteristic feature of both the components at 1150 cm**^-1^** represents the C-O bending stretching, respectively.[46] In the crosslinked HAMA1%-GelMA8% hydrogel, the characteristic peaks of gelatin and hyaluronic acid remain intact. However, the peak of C=C stretching of methacrylate group is too close to the amide I region of gelatin, making it difficult to decouple the signals for any analysis.[47] Slight variations in the intensities and position of the peaks 1500–1000 cm^-1^ suggests chemical interactions or physical blending between the HAMA1% and GelMA8% components. The described changes in the composite spectrum indicate that HAMA1% and GelMA8% interact well to form a composite hydrogel with small alterations to the chemical bonding and molecular structure compared to the individual components.

To confirm the photo-crosslinking of HAMA1%-GelMA8% hydrogels ^1^H NMR analysis was performed as shown in ***(Figure 3-I)***. The spectra were recorded at different UV irradiation times, allowing for the observation of changes as the exposure progressed. The process was evidenced by the gradual disappearance of the signals corresponding to the protons of methacryloyl vinyl (∼5.6 – 6.2 ppm) and methyl groups (∼1.9 ppm). Initially, at t = 0 s, these resonances were clearly visible, indicating the presence of unreacted methacryloyl moieties, similarly to the published GelMA and HAMA systems.[32, 39] After 45 s of UV exposure, their intensities decrease, and by 90 s, they are nearly absent. This indicates that the methacryloyl vinyl groups underwent a reaction initiated by radicals formed during photoinitiation, leading to their ‘consumption’ as they participated in the formation of new covalent bonds. The disappearance of the vinyl and methyl proton resonances confirms the ongoing reaction between the polymer chains. Furthermore, the broadening of the resonances observed in the last spectrum can be attributed to an increase in the heterogeneity of the chemical environment of the protons and the restriction of polymer chain mobility within the forming crosslinked network. The increased structural rigidity of the created three-dimensional network leads to shortening T_2_ relaxation times, contributing to peak broadening. These observations provide clear evidence of methacrylate depletion, indicating that the primary cross-linking mechanism is a radical-mediated reaction between the vinyl moieties of methacryloyl groups, leading to the formation of a crosslinked composite and corroborating with the FTIR results. The vinyl groups of the methacryloyl moieties undergo radical polymerization, forming covalent crosslinks between polymer chains (***Figure 3I, J***).

For a selected HAMA1%-GelMA8% formulation, we run both amplitude (***Figure 3K***) and frequency sweeps (***Figure 3L***) to establish their rheological properties (after crosslinking), both at room temperature (set as 22.5 °C) and at physiological temperature (37°C). Amplitude sweep demonstrated that cross-linked hydrogels largely retained their viscoelastic nature at the higher temperature, with similar storage modulus at 1% strain (***Figure 3J***). The linear viscoelastic region (LVR) was noted to be much shorter at the 37 °C, indicating the underlying gelatine chains pertain slight temperature responsiveness. Nevertheless, storage modulus still dominated over loss modulus, indicating a soft hydrogel-like structure was persistent. Frequency sweep revealed no difference between the composite HAMA1%-GelMA8% the two tested at two different temperatures (5480 ± 30 Pa at 22.5 °C and 5200 ± 290 Pa at 37 °C), taken at 1 Hz.

### 3.3. Investigation of F11 DRG Neural Cells 2D Behaviour on HAMA-GelMA Hydrogels

In this step, we investigated F11 DRG neural cell growth on the 2D surface of 500 µL hydrogel layers and analysed cell migration, morphology, viability, and the expression of neural markers. Three composite hydrogel types with varying GelMA concentrations (5%, 8%, and 10%) were tested, while the HAMA concentration remained constant at 1%. After 7 days of culture, the samples were fixed and stained for F-actin to visualize the cytoskeleton. Cell migration at HAMA1%-GelMA8% was 53.68% higher than at HAMA1%-GelMA5% and 73.81% higher than at HAMA1%-GelMA10%. The greatest, statistically significant cell migration was observed for the HAMA1%-GelMA8% hydrogels, compared to GelMA5% and GelMA10%, and therefore, they were subsequently selected for further studies (***Figure 4A, B, F***).

**Figure 4:**
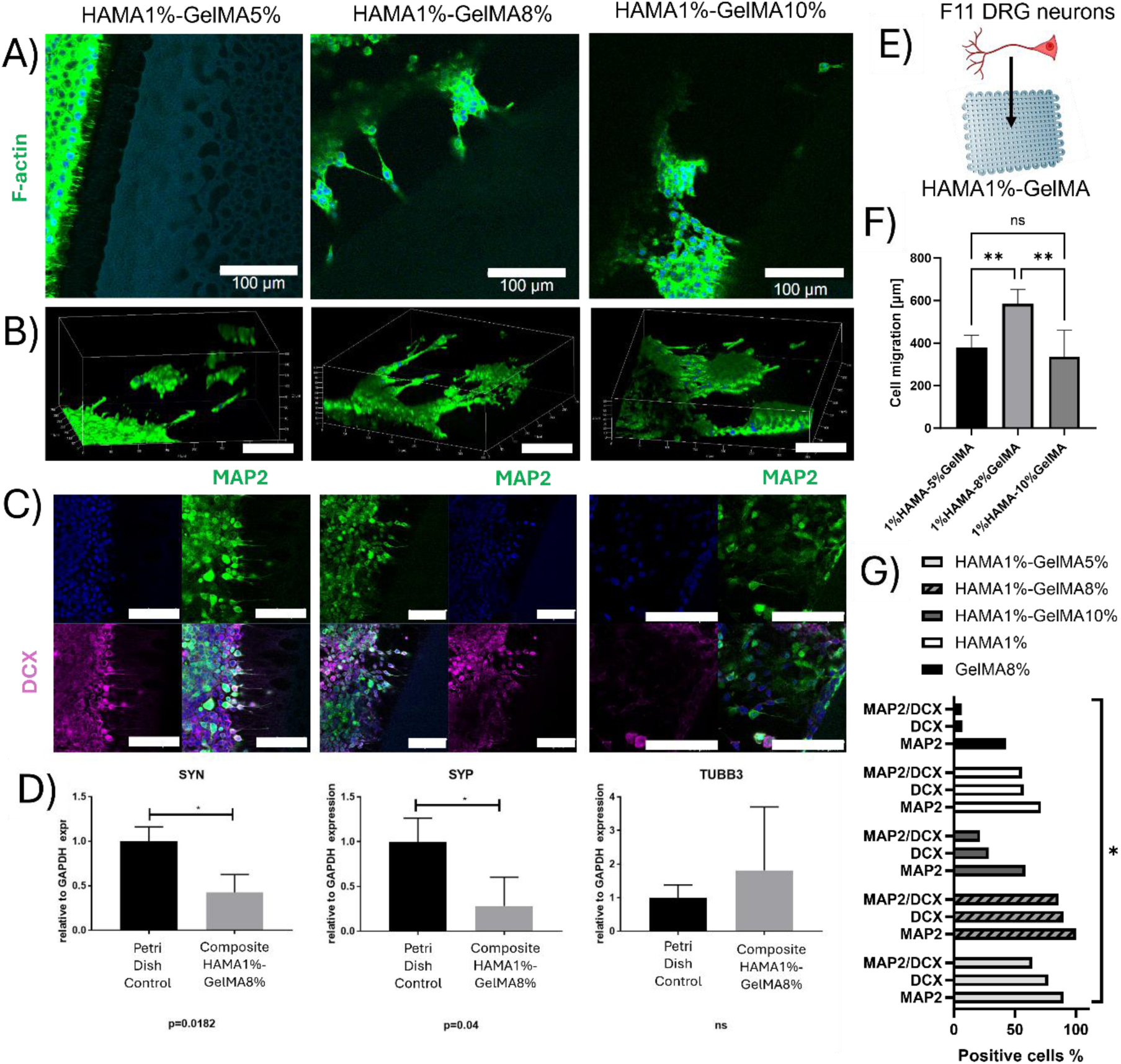
Immunocytochemical images from confocal microscopy showing F11 DRG neural cells cultured on the top of composite for 7 days A) HAMA1%-GelMA5%, HAMA1%-GelMA8%, HAMA1%-GelMA10%, stained to visualize F-actin; B) 3D reconstruction of Z-stacks; C) ICC detection of migrating (DCX, violet) and late (MAP2, green) neuronal markers in cells cultured on HAMA-GelMA composite hydrogels for 14 days. The scale bar represents 100 µm in all images. D) The differences in relative expression of the SYN, SYP and TUBB3 genes (relative to GAPDH) for F11 DRG neural cell line cultured on a top of composite hydrogel for 7 days. Control indicates cells cultured in standard Petri Dish. Statistical analysis was performed using student t-test (performed in GraphPad software) with three biological replicates. Data are presented as mean ± standard deviation; *p<0.05.

To assess the migration and maturation of DRG neural cells, immunodetection of proteins associated with migrating (doublecortin, DCX) and late (Microtubule Associated Protein 2, MAP2) neuronal markers was performed after 14 days of culture ***(Figure 4C***). The results demonstrated DCX/MAP2+ cells across all tested samples, but the most favourable cell morphology was observed on the HAMA1%-GelMA8% and HAMA1%-GelMA5%. Cell migration was not significant for the HAMA1%-GelMA5% samples, while the HAMA1%-GelMA10% hydrogels exhibited fewer DCX/MAP2+ cells in the observed sections. The results show that the composite hydrogel HAMA1%-GelMA8% demonstrated the highest percentage of MAP2/DCX positive cells, pointing towards its most effective support for neural cell migration. Specifically, 90% of cells were DCX+, while 85% were both MAP2/DCX+ (***Figure 4G***). In comparison, HAMA 1%-GelMA5% exhibited 77%, and HAMA1%-GelMA10%, 28%, significantly lower effectiveness. Interestingly, the individual control hydrogels, HAMA1% and GelMA8%, did not support DCX expression as effectively, yielding only 57% DCX+ cells for HAMA, and a mere 7% DCX+ cells for GelMA8% (***Figure S2A, B***). Cells cultured on 1% HAMA or 8% GelMA also did not exhibit proper elongated morphology and contained very few DCX/MAP2+ cells after two weeks of culture (***Figure S2A-B*** and ***Figure 4G***). These findings suggest that the HAMA1%-GelMA8% hydrogel composition is optimal for promoting the migration of F11 DRG neural cells and supporting the presence of both MAP2+ mature neurons and DCX+ migrating cells, making it the most promising candidate for further investigation (***Figure 4A, B, F***).Following systematic optimization, 8% GelMA was selected as the optimal base concentration that provided a balance between mechanical support and biological permissiveness, enabling efficient neural cell migration. Using this fixed GelMA concentration, we further investigated the effect of varying HAMA content (0.5%, 1%, 2%, and 5%) on neural cell migration, neurite outgrowth, and neural marker expression. We observed the HAMA 1%–GelMA 8% hydrogel formulation was the most favorable for promoting both neural cell migration and neurite elongation (***Figure S3A***). When using lower HAMA concentrations, such as 0.5%, we did not observe neurite outgrowth, due to insufficient mechanical integrity of the hydrogel matrix, which may degrade too rapidly or lack adequate structural support for directed neurite extension. Conversely, at higher HAMA concentrations (≥2% in combination with 8% GelMA), we observed reduced cell migration and limited neurite elongation, potentially due to increased matrix stiffness and decreased porosity, which may hinder cellular mobility and diffusion of signaling molecules. These findings are consistent with the literature, which highlights that the relationship between hyaluronic acid (HA) concentration and neural cell behavior is non-linear. Studies have shown that there is an optimal HA concentration range that supports maximal neurite outgrowth and cell motility. For example, hydrogels that are too stiff (e.g., at higher HA concentrations) can impede neurite extension due to mechanical constraints, whereas hydrogels that are too soft (e.g., at 0.5%) may degrade prematurely or lack structural cues needed for stable neurite anchorage [3, 12, 19]. Therefore, the HAMA 1%–GelMA 8% formulation appears to offer a balanced mechanical and biochemical environment that facilitates both migration and neurite elongation. To further analyse the expression of genes characteristic of neural cells, real-time qPCR was performed using primers for the markers SYP, SYN, and TUBB3. The expression of synaptic markers SYN and SYP significantly decreased in cells cultured on the composite hydrogels (p<0.05). In contrast, the expression of the TUBB3 gene, associated with neuronal cytoskeleton formation, only appeared to increase (not significant) compared to the two single component hydrogels (***Figure S2A, B***). We also performed immunocytochemical staining after 7 days in culture to detect markers of differentiated neurons, Tubulin Beta-III (TUBB3), and Synaptophysin (SYP), also known as the significant synaptic vesicle protein p38. These markers were identified in all materials we tested; however, we observed the highest number of TUBB/SYP positive cells, especially for composite hydrogel (***Figure S2C, D, E, F***). The expression of the TUBB3 gene was upregulated (not significant) at HAMA1% and HAMA-GelMA8% compared to the flat control Petri dish (***Figure S2 G***).

To further evaluate cell viability, fluorescence imaging was conducted using Calcein-AM (green) and ethidium homodimer-1 (red) after 1 and 5 days of incubation. Only live cells with proper morphology were observed, and no dead cells were detected, confirming the biocompatibility of the HAMA1%-GelMA8% hydrogel (***Figure S3***). Cell viability was assessed using flow cytometry with the Muse™ Annexin V & Dead Cell Kit. Control cells showed 98.3% survival. The results show that the cell viability of all tested materials is similar to that observed for cells cultured on the HAMA-GelMA hydrogel, suggesting that the material is not cytotoxic to F11 DRG neural cells. The comparison of cell viability across different substrates indicates that the tested material does not cause a significant decrease in live cell numbers, confirming its biocompatibility (***Figure S4***). Reactive oxygen species (ROS) were detected using the Muse® Oxidative Stress Kit. The positive control, treated with menadione, showed a significant increase in ROS-positive (ROS+) cells (approximately 60%). The lowest ROS levels were observed in cells cultured on HAMA1%, while PCL and HAMA1%-GelMA8% embedded in PCL grid (***GrooveNeuroTube***) substrates showed minimal differences. The highest ROS levels were induced in cells cultured on GelMA8% hydrogels, which confirms that the selected conditions for HAMA and the method of hydrogel preparation positively influence cell viability and do not lead to ROS generation (***Figure S5***). The elevated ROS in GelMA 8% is likely attributed to the high concentration of methacrylated gelatin and the associated photo-crosslinking process. UV-induced polymerization in the presence of photoinitiator can generate free radicals and residual oxidative species that persist in the hydrogel network and stimulate intracellular ROS production. Moreover, the high density and stiffness of 8% GelMA may limit oxygen and nutrient diffusion, further exacerbating cellular oxidative stress. The GrooveNeuroTube scaffold, incorporating this hydrogel into a PCL conduit, further decreased ROS to ∼20%, approaching levels observed for HAMA 1% alone. This progressive reduction indicates that combining GelMA with HAMA and structuring the composite within a 3D scaffold effectively mitigates oxidative stress in neural cells.

### 3.4. Impact of growth factors on cell behavior

In the next phase, we aimed to demonstrate the functionality of the hydrogels as a matrix that can further deliver a cocktail (mix) of selected growth factors. For this, we first selected several growth factors and chose their concentrations based on literature data[48–50]: NGF 2 ng/mL and 5 ng/mL, GDNF 5 ng/mL and 10 ng/mL, as well as VEGF 10 ng/mL and 50 ng/mL. In the first instance, to evaluate the general effects of growth factors on cells, we performed a simple 2D culture of F11 DRG neural cells for 7 days with the addition of these. Results revealed that the expression of neural-specific genes varied depending on the growth factor applied (***Figure S6 A-D***). As FBS is known to affect biological responses, we wanted to additionally tests its effects by modulating the concentration in the differentiating medium. In particular, the analysis showed that DCX expression was significantly different between cells cultured in 1% FBS (differentiating medium) and those in 10% FBS (standard culture medium). The treatment of 2 ng/mL NGF notably increased DCX expression in the cells, while 5 ng/mL NGF had an inhibitory effect (***Figure S6A***). The use of the other GF (GDNF, VEGF) resulted in higher DCX expression compared to standard culture conditions (***Figure 5A***). The increase in peripherin expression was observed with NGF at 2 ng/mL, GDNF at 10 ng/mL, and VEGF at 50 ng/mL (***Figure S6B***). Treatment with 2 ng/mL NGF and 50 ng/mL VEGF stimulated peripherin expression relative to 1% FBS culture (without statistical significance) and standard culture with 10% FBS (statistically significant differences) (***Figure S6B***). For SYN and S100b, increased expression was observed in samples treated with NGF and 50 ng/mL VEGF and 2 ng/mL, respectively (***Figure S6C, D***). Subsequently, to assess morphological changes, imaging of cells cultured for 7 and 14 days was captured (***Figure 6F***). In those experiments, we tested NGF (2 ng/mL and 10 ng/mL), GDNF (5 ng/mL and 10 ng/mL), and VEGF (10 ng/mL and 50 ng/mL). Given the noted individual effects of growth factors, we generated two selections of their mixtures: **MIX1** contained 2 ng/mL NGF, 5 ng/mL GDNF, and 10 ng/mL VEGF, while **MIX2** contained 10 ng/mL NGF, 10 ng/mL GDNF, and 50 ng/mL VEGF. After 7 days of incubation, cells were collected by trypsinization, and proliferation and viability were assessed using Trypan Blue staining. Results are summarized in ***Figure S6F-H.*** The results demonstrate that cell numbers remained consistently high across all conditions over the 72-hour culture period. Notably, the highest proliferation rates were observed in samples treated with **MIX1**, reaching up to 3.3 million cells, suggesting that this combination provided the most favourable environment for cell growth. Additionally, treatments with GDNF 10 ng/mL, VEGF 50 ng/mL, and **MIX2** also promoted significantly higher cell numbers than the control, which maintained a stable, albeit a lower count throughout the experiment. Importantly, prolonged culture did not result in a decline in cell numbers, indicating that the experimental conditions effectively supported sustained cell viability and proliferation (***Fig S6 F, G, H***).

**Figure 5:**
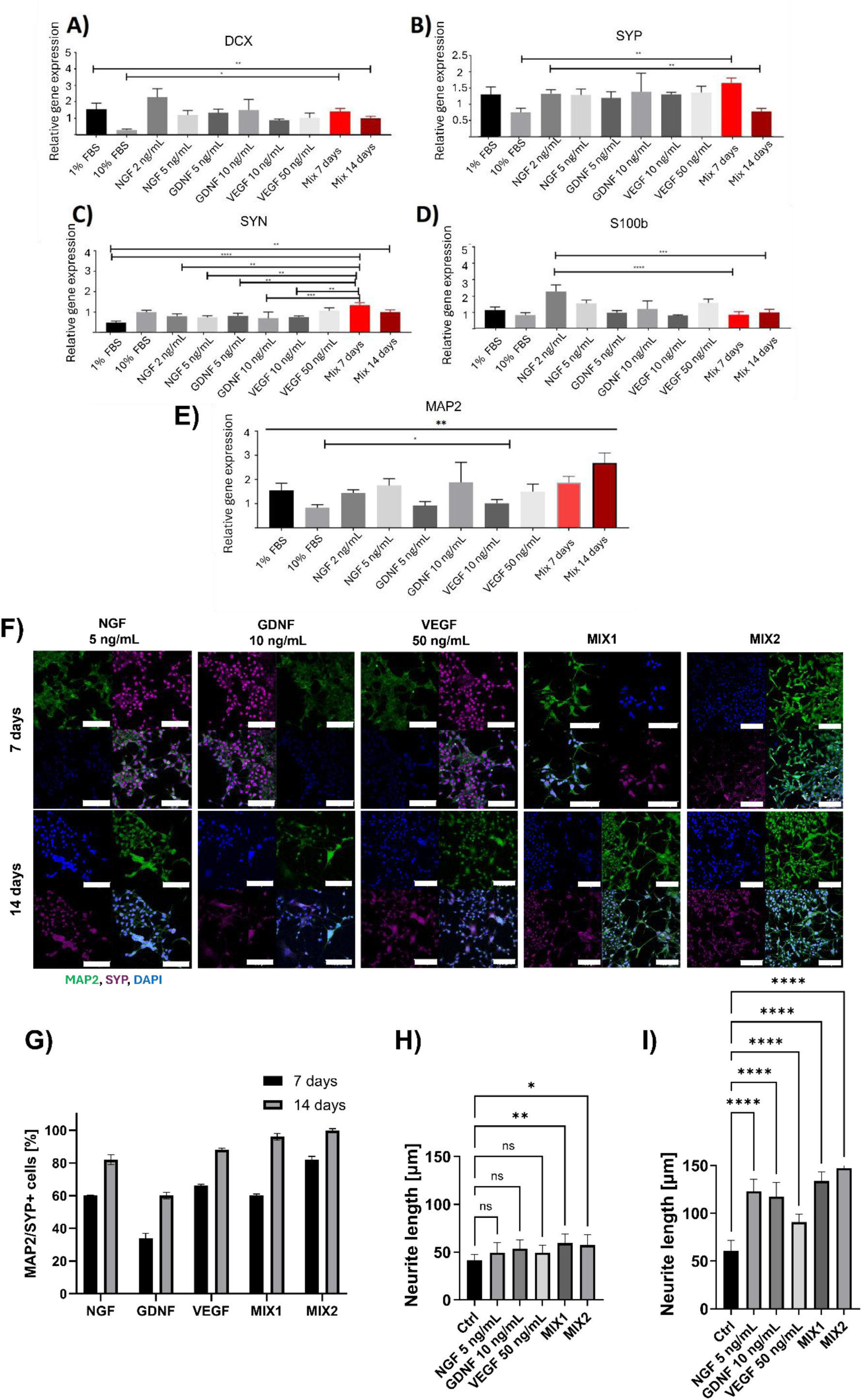
The differences in the relative expression of the A) DCX, B) SYP, C) SYN, D) S100b, and E) MAP2 genes (relative to GAPDH) for F11 DRG neural cell line cultured in the presence of growth factors for 7 and 14 days. F) Visualization of F11 DRG neural cells subjected to growth factors for 7 and 14 days. Representative confocal images immunostained for MAP2 (mature neurons; green) and SYP (synaptic marker, violet) (bottom panel) cultured for 21 days in the presence of different growth factors and their mixes. The scale bar represents 100 µm in all images. G) The histograms show the quantification of positive to total number of cells corresponding to MAP2/SYP+. Neurite length after culture of H) 7 days and I) 14 days with GFs. Statistical analysis was performed using One-Way ANOVA (performed in GraphPad software) with Tukey’s post hoc test. Data are presented as mean ± standard deviation; n=3, *p<0.05, **p<0.01, ***p<0.001, ****p<0.0001.

**Figure 6:**
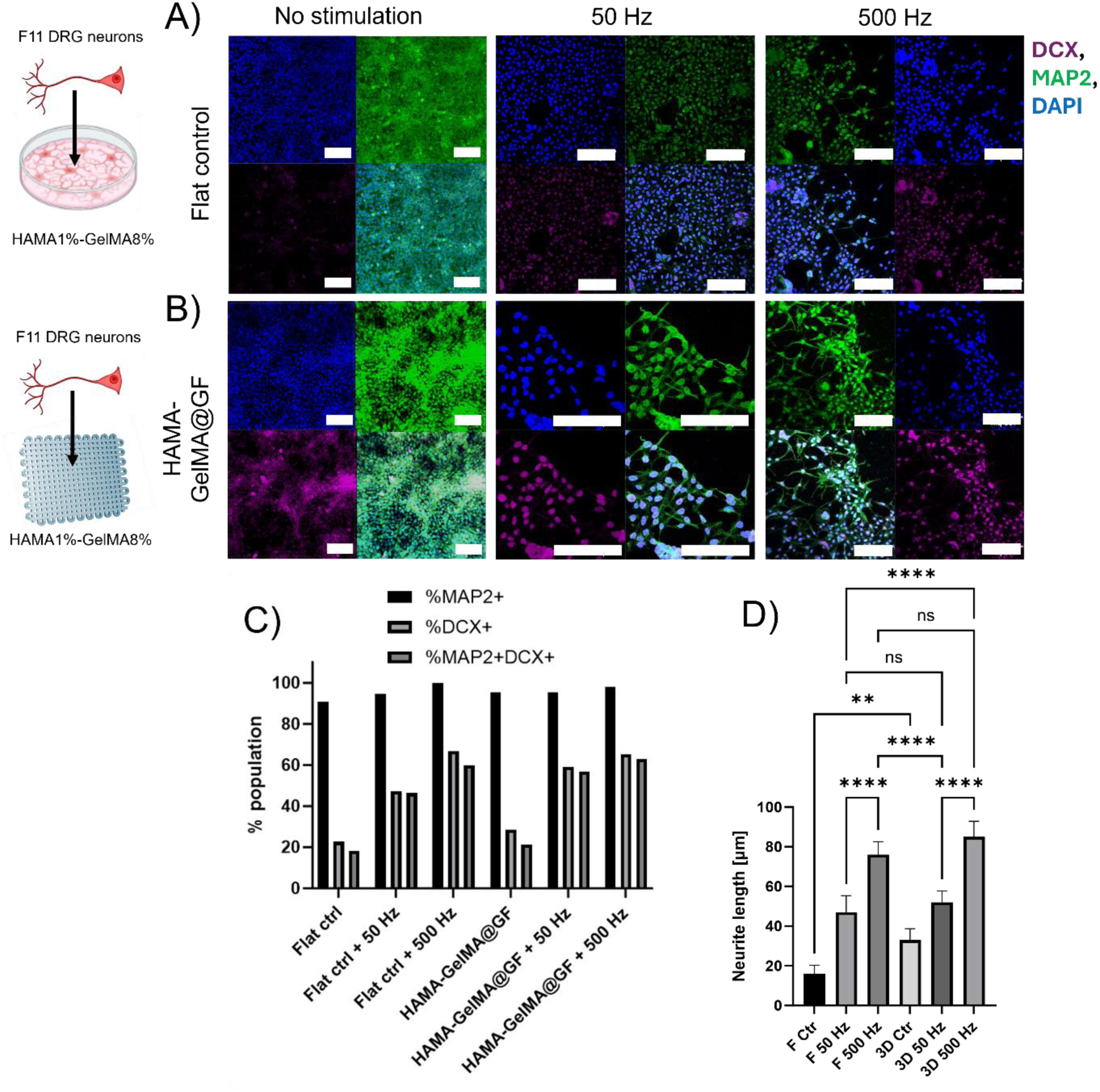
Representative confocal images of F11 DRG neural cells cultured on A) flat control Lab-Tek and B) HAMA1%-GelMA8%-GF(MIX2) cultured in control conditions and subjected to 50 Hz or 500 Hz PEMF. Cells were immunostained for MAP2 (mature neurons; green) and DCX (a marker of migrating neurons, violet). Cells were exposed to AC at 50 Hz and 500 Hz for 3 days. The scale bar represents 100 µm in all images. C) The histograms show the % population of MAP2, DCX and double MAP2/DCX positive cells at various frequencies, D) Neurite length of F11 DRG neural cells. Statistical analysis was performed using One-Way ANOVA (performed in GraphPad software) with three biological replicates. Data are presented as mean ± standard deviation; *p<0.05, **p<0.01, ***p<0.001, ****p<0.0001.

F11 DRG neural cells displayed noticeable differences in morphology. The most prominent observation was the elongation of neurites in cells treated with MIX1 and MIX2. Images of F11 DRG neural cell morphology were captured after 7 and 14 days of culture using an optical microscope (***Figure S6E***). It was noted that the growth factors influenced significant differences in cell morphology. Elongated morphology and long neurites were observed in both MIX-treated groups, suggesting the progression of neural maturation (***Figure S6E***).

The expression of the DCX gene in the samples after 14 days of culture significantly increased in the 7-day culture compared to the 10% FBS control group but decreased in the 14-day culture when compared to the 1% FBS group (***Figure 5A***). For the SYP gene expression, a significant statistical increase was observed in the 7-day culture with the growth factor mix compared to the 10% FBS control group, while a significant decrease was noted in the 14-day culture compared to the group with 1% FBS and 2 ng/mL NGF (***Figure 5B***). Regarding the SYN gene expression (***Figure 5C***), a statistically significant increase was observed after 7 days of culture in the presence of 1% FBS and the growth factor mix, compared to almost all other samples (except the 10% FBS control). The expression of the 100b gene significantly decreased in cells cultured with 1% FBS and a growth factor mix for both 7 and 14 days, compared to cells cultured with 1% FBS supplemented with 1% FBS and 2 ng/mL NGF (***Figure 5D***). Analysis of MAP2 expression (***Figure 5E***) indicated a statistically significant increase in expression in cultures with 1% FBS and 10 ng/mL GDNF compared to the 10% FBS control group. After 14 days of culture, expression levels were further increased.

Immunocytochemical (ICC) detection of markers typical for mature neurons (MAP2) and synapses (SYP) revealed differences in the number of MAP2/SYP+ positive cells, depending on the growth factors used (***Figure 5F***). After 7 days of treatment with growth factors, the highest percentage of MAP2/SYP+ cells was observed with VEGF (66%), and the lowest with GDNF (44%). For MIX1 and MIX2, the results were 60% and 82%, respectively (***Figure 5G***). After 14 days, all groups showed an increase in the percentage of positive cells, with MIX1 and MIX2 still achieving the highest values—96% and 100%, representing an increase of 36% and 18%, respectively, compared to day 7 (***Figure 5G***). When comparing both time points, the largest increase in the percentage of positive cells was seen in the MIX1 (36% increase) and MIX2 (18% increase) groups, which correlates with the PCR results. The analysis of neurite length in cells treated with various growth factors over 7 and 14 days reveals significant differences, particularly with MIX1 and MIX2. After 7 days, the average neurite length was highest in the MIX1 group (59.3 µm), followed closely by MIX2 (57.3 µm), while the control and other growth factors like NGF and GDNF showed lower average lengths, ranging from 41.4 µm to 53.7 µm (***Figure 5H***). After 14 days, MIX1 and MIX2 exhibited further growth, with MIX2 showing the most extended neurites (154.3 µm) compared to other groups (***Figure 5I***). The average neurite length in the MIX1 group was also significantly higher (133.6 µm) compared to the control (60.7 µm) and other growth factors like VEGF and NGF. These results indicate that MIX1 and MIX2 promote substantial neurite extension, with MIX2 demonstrating the most pronounced effect over the 14 days (***Figure 5I***). From these results, we concluded that MIX2 (containing 10 ng/mL NGF, 10 ng/mL GDNF, and 50 ng/mL VEGF.) provides the most suitable migration and neurite outgrowth of the F11 DRG neural cells, and hence fixed it for further studies.

### 3.5. Morphological changes of F11 DRG neural cells under PEMF

Non-invasive therapies like PEMF offer exciting potential for advancing nerve repair strategies. PEMF has been shown to promote neurite extension, enhance neural cell viability, and reduce inflammation, positioning it as a powerful adjunct to material-based interventions. The combination of these approaches seems to address current limitations and offer ways to achieve superior outcomes in nerve repair.[34] Before testing developed ***GrooveNeuroTube***, we sought to find the optimal parameters for applying PEMF. We first conducted PEMF stimulation for 3 days on flat control substrates (standard Lab-Teks substrates) and cells cultured on hydrogels using an in-house PEMF stimulation system (***Figure S7***). We chose 50 and 500 Hz sinusoidal shape functions for experiments, providing a 0.24 mT alternating magnetic field induction in the spectrum used across medical applications.[51, 52]

On day 3 of the culture, we performed ICC to detect the expression of DCX and MAP2 on both control substrates as well as HAMA1%-GelMA8% composite hydrogel containing MIX2 of growth factors (named as HAMA1%-GelMA8%-GF) (***Figure 6 A, B***). The data shows that exposure to PEMF (both 50 Hz and 500 Hz) significantly influences neural cell markers (MAP2, DCX, and MAP2/DCX) in both 2D (Flat) and 3D (HAMA-GelMA@GF) cultures. In the 2D system, PEMF 500 Hz treatment led to a percentage increase in MAP2-positive cells from 94.7% to 100%, compared to PEMF 50 Hz (***Figure 6C***). Similarly, DCX-positive cells increased from 47.4% to 66.7% (***Figure 6 A, C***). In the 3D HAMA-GelMA@GF environment, the response to PEMF treatment was more pronounced. PEMF 500 Hz resulted in in an increase in MAP2+ cells from 95.5% to 97.8%, and DCX+ cells from 59.1% to 65.2%compared to PEMF 50 Hz (***Figure 6 B, C***). These findings suggest that the synergistic combination of the 3D hydrogel scaffold and PEMF, especially at 500 Hz, significantly enhances neural marker expression compared to 2D conditions, indicating improved neural differentiation and migration. The highest percentage of double-positive MAP2+DCX+ cells was found in the HAMA-GelMA@GF + 500 Hz condition (63.04%), further supporting the enhanced neurogenic commitment under this condition. More pronouncedly elongated MAP2/DCX+ cells were observed under the PEMF treatment condition (***Figure 6 A, B***). The highest number of cells positive for both DCX and MAP2 was found at 500 Hz on Lab-Tek controls and flat hydrogels (***Figure 6C***). Elongated cells with long neurites were observed on hydrogels subjected to 50 and 500 Hz. The most extended neurites were found at 500 Hz, with neurites reaching over 80 µm in length (***Figure 6D***). We also confirmed the cell viability after culture and treatment with PEMF at two selected frequencies (50 and 500 Hz) for 24 hours. Cells were cultured on 96-well plates, with substrates including HAMA, GelMA, HAMA1%-GelMA8%, and PCL-HAMA1%-GelMA8% (unrolled ***GrooveNeuroTube***). The PEMF stimulation did not negatively affect cell viability, with an average of 10% more cells showing improved viability after PEMF stimulation than those cultured without stimulation (***Figure S8***). We chose the square 500 Hz condition for further experiments based on the data because it demonstrated the highest cell viability, particularly in the PCL-HAMA-GelMA group, where cell viability reached 99.26 ± 0.9% at 500 Hz. In comparison, the viability for PCL-HAMA-GelMA at 50 Hz was significantly lower, at 74.1 ± 0.66%. This marked improvement in cell viability at 500 Hz made it the optimal experimental condition for further studies. From these preliminary tests, we selected this stimulation to study our ***GrooveNeuroTube ex-vivo*** model over a prolonged 60-day period in the final stages.

### 3.6. Translation of 3D cell-laden collagen hydrogel in *GrooveNeuroTube* toward better *in vivo* resemblance

To evaluate the ***GrooveNeuroTube*** (containing HAMA1%-GelMA8%-GF), we performed a 3D bioprinting of F11 DRG neural cells on both ends of the tube to simulate *ex vivo*-like conditions (***Figure 1C***). Experiments were conducted under standard culture conditions and under conditions subjected to PEMF stimulation, selected as 500 Hz, as described in the previous section. Cross-sections of the tube were made by vibratome cutting, which allowed for accurate sectioning of 500 µm slices of the desired areas on days 1, 14, and 28 of culture. It can be observed that at the corner of the gel after 1 day of culture (***Figure 7A***, arrow pointing at a schematic location of the tube), according to the design, there is a clear distinction between cells labelled in the collagen hydrogel versus empty area of the tube. The distributions of cell orientation angles indicate random directional growth on day 1 (***Figure 7B***). In the central part of the tube (black arrows guiding locations in ***Figure 7G***), after 14 and 28 days, living cells labelled with Calcein-AM are visible, and their numbers increased with each week of culture (***Figure 7H***). The cells are evenly and randomly ***(Figure 7D***) distributed within the hydrogel in the 14 days of culture. In the case of the PEMF-treated group, we observed more cells within the central location, and they were aligned in the direction of the tube compared to the cells without PEMF (***Figure 7F***). After 28 days, a clear presence of many live cells and a high ratio of live-to-dead cells were observed (***Figure 7H***). In the case of ***GrooveNeuroTubes*** cultured for 28 days without PEMF, the cells showed only slight organization within the hydrogel (***Figure 7D***). However, for samples subjected to PEMF, the cells were organized along the tube exposed to PEMF (***Figure 7F***). Cell viability results showed that, on day 1, 71.72 ± 0.28% of the cells were live, increasing to 86.67 ± 0.33% by day 14 (***Figure 7H***). After PEMF exposure on day 14, live cells reached 91.21 ± 0.08%, and by day 28, 92.02 ± 0.07% of the cells were alive. For samples exposed to PEMF for 28 days, live cell viability improved further to 95.05 ± 0.05%, significantly reducing dead cells (4.95% ± 0.05%) (***Figure 7H***). We could conclude that up to 28 days of culture, the PEMF stimulation already shows beneficial signs for the growth of neural cells. To determine whether cells seeded from both ends of the GrooveNeuroTube migrated and converged in the central region after 28 days in culture, we performed differential fluorescent cell labeling at each end of the conduit. Specifically, cells seeded at one end were labeled with CellTrace™ Far Red DDAO-SE, enabling detection in the far-red channel (***Figure 7J***), while cells seeded at the opposite end were labeled with CellTrace™ CFSE, which is excited at 488 nm and detected in the green channel (emission ∼517 nm) (***Figure 7K***). After 28 days of culture, fluorescence imaging revealed clear co-localization of both cell populations in the central part of the conduit, indicating successful bidirectional migration and convergence (***Figure 7L-M***). Cells from both ends migrated approximately 8.25 mm each, meeting at the center of the ∼16.5 mm-long tube. This experiment confirms symmetric migration dynamics and complete bridging of the conduit by day 28. In the central region of the GrooveNeuroTube, differences could be observed between the PEMF-stimulated and non PEMF-stimulated conditions. In particular, a higher density of cells was detected under PEMF stimulation, enhancement of proliferation and survival (***Figure 6I***). These findings align with previous reports showing that pulsed electromagnetic fields (PEMF) promote both neurite outgrowth and proliferation in neuronal-like cell lines, including F11 cells, which are derived from dorsal root ganglia. In our system, we also observed that, consistent with previous findings in other experimental conditions, cells exhibited longitudinal alignment along the axis of the conduit, a phenomenon further supported by the presence of directional neurite extension (***Figure 6M***). This alignment is likely influenced by both the architectural guidance of the mesh structure and the directional cues enhanced by PEMF exposure.

**Figure 7:**
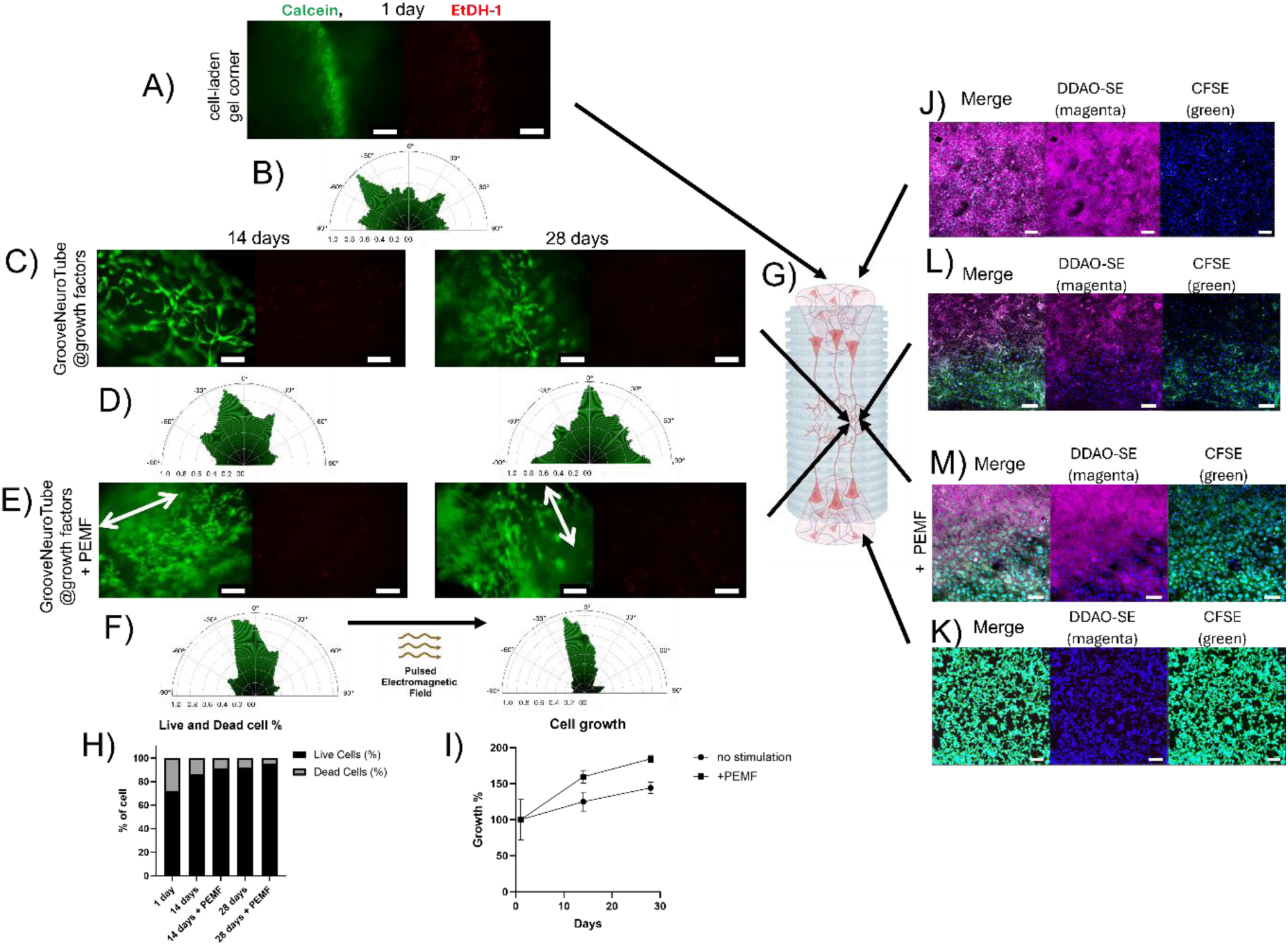
Confocal images of live/cell assay of live (Calcein-AM labelled; green) and dead (EtDH-1, red) neural cells cultured in 3D bioprinted tubes at the ends of the **GrooveNeuroTube** for A) 1, B) 14 and D) 28 days. Cells were exposed to PEMF magnet stimulation according to the parameters selected in previous experiments (PEMF) = AC 500 Hz. B, D, F) Circular histograms for the orientation of the cells direction, after culturing for 1, 14 and 28 days in **GrooveNeuroTube** cultured in standard conditions C) and E) subjected to PEMF. G) Scheme of the cross-section place, H) % of live and dead cells, I) % of cell growth with and without PEMF; J–M) Confocal images of DRG neurons bioprinted in collagen-mTG within the GrooveNeuroTube and labeled with fluorescent cell trackers. J) Neurons on the upper side of the tube labeled with CellTrace™ Far Red DDAO-SE (magenta). K) Neurons on the lower side of the tube labeled with CellTrace™ CFSE (green). L–M) Cross-sectional views of the middle part of the tube after 28 days in culture: L) without PEMF stimulation and M) with PEMF stimulation. Channels are shown individually; the absence of signal in one channel reflects selective labeling of cells with a single fluorophore. The scale bar represents 100 µm in all panels. The scale bar represents 100 µm in all images.

Next, to assess whether cell migration is influenced solely by PEMF, we tested our ***GrooveNeuroTube*** again with or without the addition of growth factors (***Figure 8***). These were tested over a period of 1 to 60 days under long-term conditions to stimulate *ex vivo* experiments and reduce *in vivo* testing on animals. The results revealed that in all conditions, F11 DRG neural cells were able to migrate to the centre of the tube over a period of 28-60 days. This was confirmed by performing transverse cross-sections at 28 and 60 days, followed by ICC testing, which identified the cells (***Figure 7C, E*** and ***8A, B, C, F, G***). We observed elongated cells and multicellular clusters migrating from the cell-laden hydrogel segments that were 3D bioprinted at both ends of the tubular scaffold. Migration distance was determined by measuring the extent of these advancing clusters, which emerged from the hydrogel regions and progressed along the inner surface of the tube toward the central region. The most remarkable cell migration was noted for the ***GrooveNeuroTube*** containing growth factors and under PEMF stimulation (**Figure 8C, G**). The data reveals that cell migration varied depending on the conditions applied, mainly when active agents and PEMF were used. The migration distance in the GrooveNeuroTube without growth factors group ranged from 0.074 mm on day 1 to 3.52 mm on day 60 (***Figure 8G***). The ***GrooveNeuroTube+GF*** group showed higher migration, with distances starting at 0.1064 mm on day 1 and reaching 5.46 mm by day 60 (***Figure 8G***). The ***GrooveNeuroTube+GF+PEMF*** group demonstrated the highest migration, with cells migrating 0.1 mm on day 1 and extending up to 7.2 mm on day 60. For the ***GrooveNeuroTube+GF+PEMF*** group, the average migration across all time points was 3.7926 ± 1.3039 mm. The highest observed migration was 7.2 mm at day 60, representing an impressive 88% greater migration than the ***GrooveNeuroTube+GF*** group, and 106% more than the ***GrooveNeuroTube*** group. On average, cells in the ***GrooveNeuroTube+GF+PEMF*** group migrated 3.79 ± 1.30 mm, which is a notable improvement compared to the 2.72 ± 1.46 mm in the ***GrooveNeuroTube+GF*** group and 1.39 ± 0.93 mm in the ***GrooveNeuroTube*** group. These results indicate that PEMF, in combination with active agents, significantly enhanced cell migration (***Figure 8G***). On day 60, most of the observed cells were dispersed throughout the entire volume of the construct, indicating that they had colonized nearly the whole structure. To confirm this, an analysis of the area covered by cells was performed. The highest cell density on day 60 was indeed observed in the PEMF-treated group (***Figure 8C***). The results demonstrate that the ***GrooveNeuroTube+GF+PEMF*** exhibited the highest cell coverage area at all time points, showing a marked increase over the duration of the experiment. Starting at a relatively low level on day 1, the cell coverage in this group grew significantly, reaching 95 ± 7% on day 60, representing an increase of approximately 850%. This suggests combining growth factors and PEMF greatly enhances cell migration over time (***Figure 8C***). The ***GrooveNeuroTube+GF*** group also showed a steady increase in cell coverage, starting at 7-10 on day 1 and rising to 88 ± 6 % by day 60, corresponding to a 700% increase (***Figure 8B***). This indicates that growth factors alone have a considerable impact on promoting cell migration, although the effect is less pronounced compared to the group with PEMF. In contrast, the PCL+GelMA tube - No HAMA@GF control group consistently demonstrated the lowest cell coverage area across all time points, indicating the importance of both components in composite hydrogel for the therapeutical effect. Initially, this group started with a cell coverage of 6-12 on day 1, and although there was some growth over time, it only reached 39 ±4 % by day 60, which represents a 550% increase (***Figure 8A***). This lower growth rate suggests that the absence of growth factors and PEMF results in a less efficient cell migration process. In summary, the data clearly shows that the combination of growth factors + PEMF leads to the most significant and sustained increase in cell coverage, indicating a potent synergistic effect on cell migration. Meanwhile, the growth factors alone have a notable but comparatively more minor impact, and the PCL+GelMA tube - No HAMA@GF group demonstrated the least response, highlighting the importance of both growth factors and PEMF in promoting cell migration (***Figure 8H***). This proved that the F11 DRG neural cells could freely migrate within the developed hydrogel composition and that the additional PEMF stimulation enhanced their migration.

**Figure 8:**
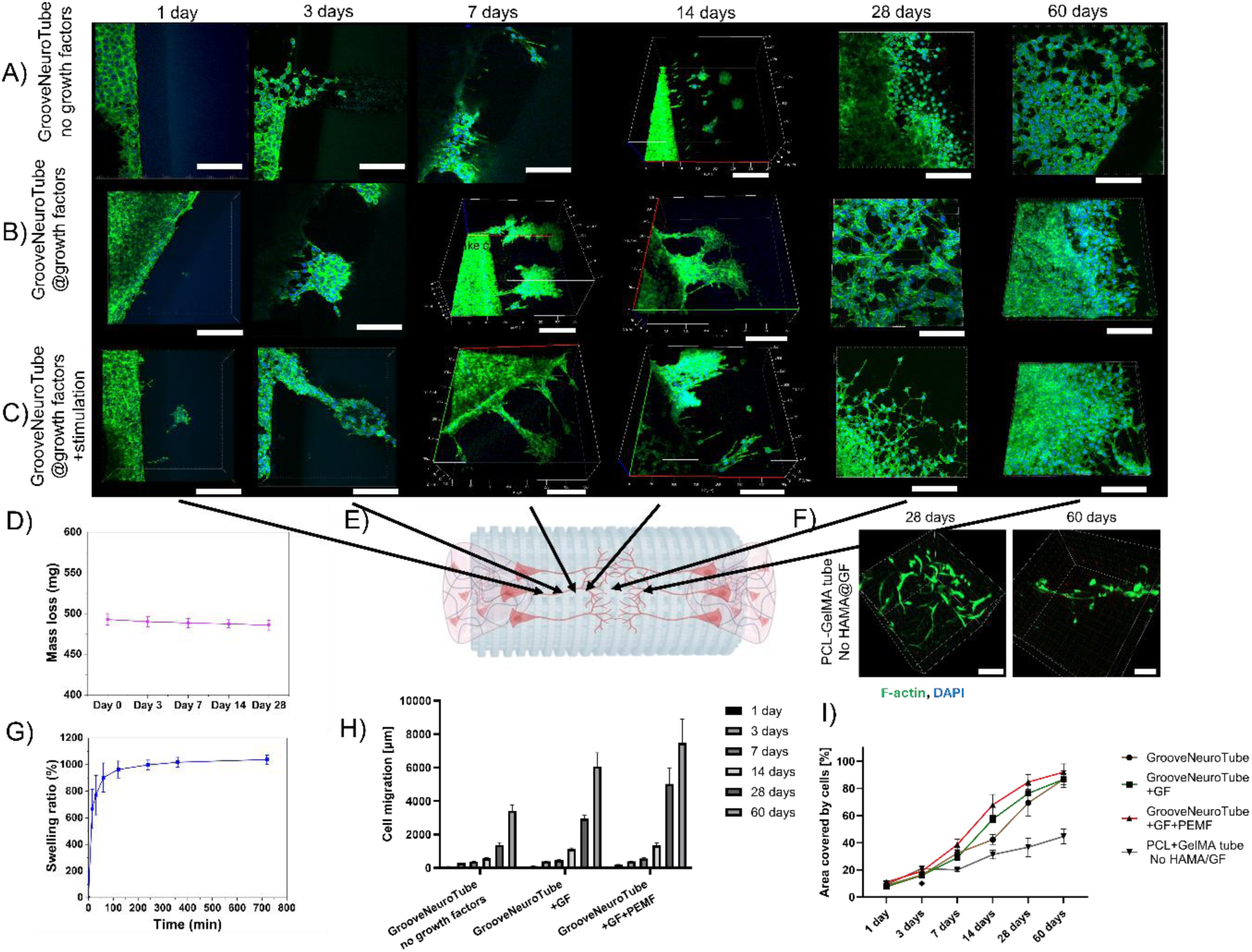
Representative confocal images of stained F-actin cytoskeleton (green, Pallodin) and nucleus (blue, DAPI) in F11 DRG neural cells 3D bioprinted at the end of the: A) HAMA1%-GelMA8% without growth factors, B) HAMA1%-GelMA8% with growth factors (**GrooveNeuroTube**), C) HAMA1%-GelMA8% with growth factors (**GrooveNeuroTube**) + PEMF, D) Degradation of GrooveNeuroTube, E) Schematic representation of the crossection, F) Tube made of only PCL and GelMA8% as control, after 28 and 60 days in culture stained for F-actin cytoskeleton (green, Phalloidin) and nucleus (blue, DAPI). The scale bar in each image represents 100 µm., G) Swelling ratio, H) Cell migration analysis, I) Total area covered by cells. The scale bar represents 100 µm in all images.

The swelling ratio of HAMA1%-GelMA8% hydrogel shown in ***Figure 8G*** indicate absorption 1̴ 000% its dry weight indicating high water retention capability. The degradation (***Figure 8D***) of the GrooveNeuroTube shows steady and minimal mass loss of 2̴ % over 60 days indicates excellent structural stability under physiological environment. The PCL microfibers ensure long-term stability, while the hydrogel enhances bioactivity and supports cell migration and axonal regrowth. Unlike solid-walled PCL-based conduits with slow degradation [1,3,4], our fibrous architecture offers controlled support without long-term tissue persistence.

### 3.7. Characterization of neurites

To confirm the functional process of neuronal maturation, we performed ICC to determine the presence of MAP2 and SYP proteins, characteristic of synaptic vesicles, which are considered synaptic markers, over 60 days of culture. After 7 days of culture, a significant proportion of the MAP2/SYP+ cells with an elongated shape were present on the PEMF-treated tube, compared to the sample without growth factors and PEMF (***Figure 9***). However, starting from day 7, highly elongated MAP2/SYP+ cells, closely adjacent to each other, with a strong SYP fluorescence signal found at their intercellular junctions, were observed after 7, 14, 28, and 60 days of culturing under PEMF conditions. These cells were kept nearby, presumably allowing inter-cellular connections. Cells cultured within the tube without active agents and PEMF surfaces (***Figure 9A***) were DCX/MAP2/SYP+. However, the cells’ shape was not polarized and did not resemble those grown on PEMF, as their neurites were shorter than the cells cultured (***Figure 9B***). The results of the analysis show a clear impact of growth factors (GF) and PEMF stimulation on the expression of neuronal markers, particularly MAP2, SYP, and their co-expression (MAP2/SYP), over time. In conditions without growth factors (-GF), the percentage of MAP2-positive cells increases steadily from 60% at 7 days to nearly 100% by 60 days, suggesting progressive neuronal differentiation without external growth signals (***Figure 9 E***). However, the expression of SYP, a marker for synaptic proteins, is considerably lower under -GF conditions, starting at 20% on day 7 and fluctuating across time points, peaking at 86.67% on day 28 before slightly declining to 66.67% on day 60. The co-expression of MAP2/SYP in -GF conditions is also relatively low, with an initial value of 15% on day 7, rising to 83.33% by day 28, but dropping back to 43.86% by day 60. In contrast, the conditions with growth factors and PEMF (***GrooveNeuroTube+GF+PEMF***) (***Figure 9B***) led to a marked increase in the percentage of MAP2-positive cells throughout the experiment. Starting at 90.91% on day 7, the percentage continued to increase, reaching 98.63% by day 60. This suggests that combining GF and PEMF significantly enhances neuronal differentiation and maturation. SYP expression in these conditions was also significantly higher than in the -GF group, beginning at 78.79% on day 7, peaking at 88.46% by day 14, and stabilizing around 87.67% at day 60. The MAP2/SYP co-expression followed a similar trend, starting at 72.73% on day 7, rising to 76.92% by day 14, and reaching 82.19% by day 60, indicating that GF and PEMF treatment promotes not only neuronal differentiation but also synaptic maturation over time ***(Figure 9 E).*** The data reveals a time-dependent effect of growth factors and PEMF on neuronal marker expression. While MAP2 expression continues to increase in both the -GF and GF+PEMF conditions, growth factors and PEMF consistently result in higher levels of MAP2, SYP, and MAP2/SYP co-expression, especially at later time points. In particular, the GF+PEMF condition enhanced synaptic protein expression (SYP) and neuronal maturation (MAP2/SYP co-expression) more effectively than -GF conditions, demonstrating the combined positive impact of growth factors and PEMF on neuronal development and synaptic protein expression.

**Figure 9:**
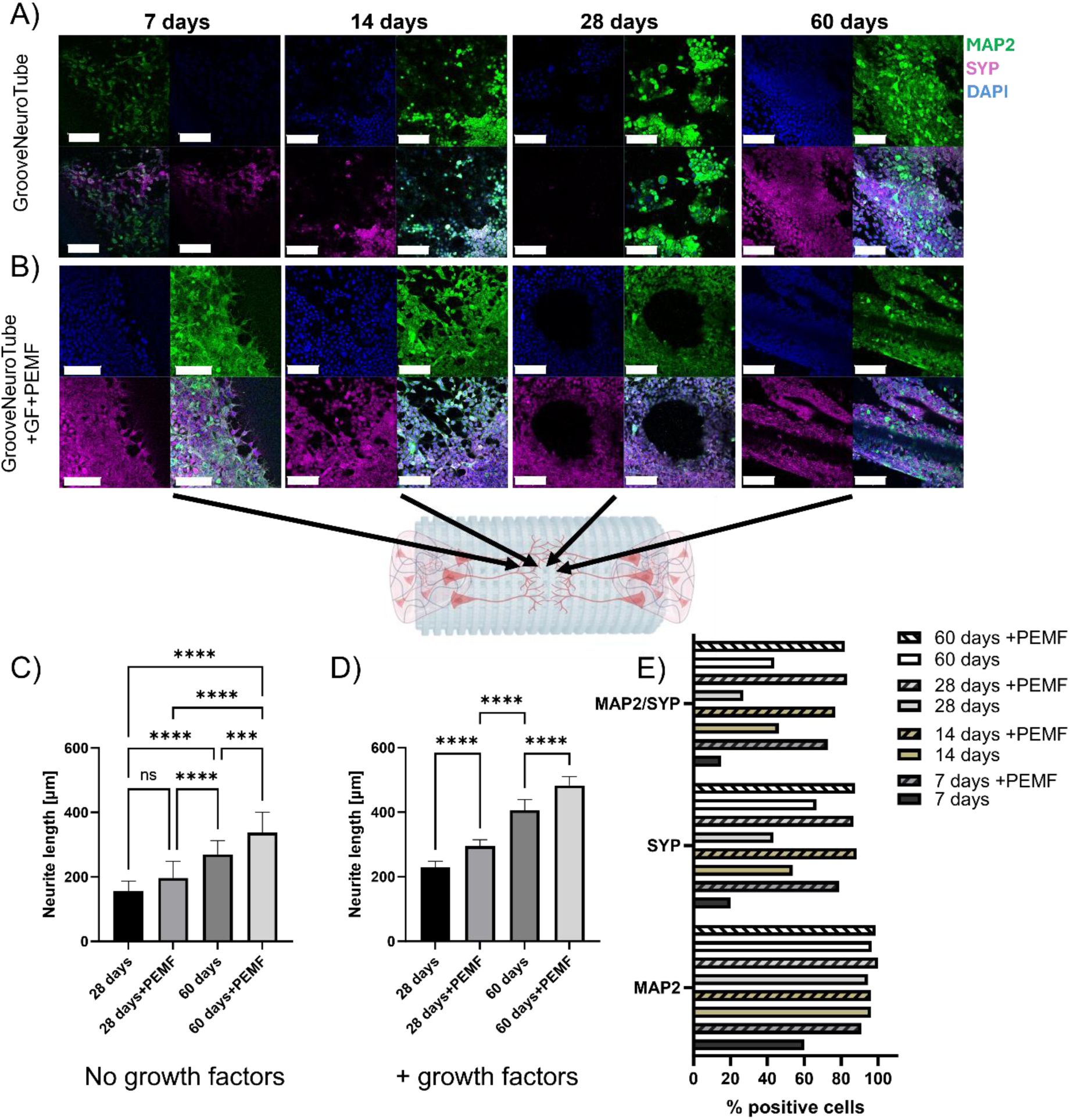
Confocal images of F11 DRG neural cells under 7, 14, 28, and 60 days in culture, immunostained for MAP2 (mature neurons; green) and SYP (synaptic marker, violet). A) The upper panel shows cell orientation within the **GrooveNeuroTube+GF** and B) The bottom panel presents cell orientation after exposure to PEMF (**GrooveNeuroTube+GF+PEMF**). The scale bar represents 100 µm in all images. C) Mean neurite length of neural cells cultured for 28 and 60 days within a tube without growth factors, D) Mean neurite length cultured for 28 and 60 days within a tube with growth factors. For both C and D cases with and without PEMF stimulation are presented. The population means (represented by each bar) essentially differ at the significance level - 0.01. E) The histograms show the quantification of the number of positive /total number of cells corresponding to MAP2/SYP+; the asterisk indicates significant differences at p < 0.01; ****p<0.0001. ANOVA

Next, we analysed the neurites’ outgrowth of F11 DRG neural cells within the tube for 28 and 60 days with conditions without and with PEMF (***Figure 9C, D***). Consecutively, we examined the average length of the neurites. With and without stimulation, DRG neural cells cultured within ***GrooveNeuroTube*** showed significant differences in neurite length (***Figure 9C***). The average length of neurites in ***GrooveNeuroTube*** without active agents and without PEMF after 28 days was 156 ± 29 µm, whereas in the PEMF-treated sample, the average neurite length was 196 ± 49 µm. The average length of neurites after 28 days within the tube with growth factors and PEMF was 229 ± 18 µm and 295 ± 19 µm in the tube with PEMF. On the 60th day, it was 406 ± 32 µm in the ***GrooveNeuroTube*** with active agents without PEMF and 482 ± 27 µm in the ***GrooveNeuroTube*** with PEMF (***Figure 9C, D***).

## 3.8. Antibacterial effects of GrooveNeuroTube

To finally demonstrate another possibility for future studies on our ***GrooveNeuroTube***, we briefly demonstrate the encapsulation of lysozyme, a known antimicrobial agent.[53] We studied its effects over a more extended period of time (up to 21 days) against *S. aureus* and *E. coli*. To do so, we loaded lysozyme within HAMA1%-GelMA8% hydrogel and examined bacterial inhibition over 7, 14, and 21 days. Detailed results are described in the supplementary materials. The lysozyme solution alone (Lyz 0.5 mg/mL) demonstrated moderate antibacterial effects. Still, the GelMA-HAMA-Lyz hydrogel was more effective in reducing bacterial growth over time to inhibit over 80% of the bacteria within 21 days (***Figure S9***). These results indicate the sustained antibacterial potential of the GelMA-HAMA-Lyz hydrogel against *E. coli* (***Figure S9D***). The F11 neural cells treated with lysozyme for 24 (***Figure S10***), 48 (***Figure S11***), and 72 (***Figure S12***) hours showed no evident toxic effect on F11 DRG neural cells. Cell morphology under each condition was compared to the untreated control conditions, and no significant morphological changes were observed, indicating that lysozyme treatment did not induce toxicity at any of the time points analysed. This confirms that the ***GrooveNeuroTube*** construct can be used to study the effects of antibacterial agents and mimic infection-like scenarios in future studies.

## 4. Conclusions

This study aimed to demonstrate the practicality of using PCL grid substrates with internal fibers to create tubes with topographical cues. This structure protects the composite hydrogel we developed, containing HAMA1%-GelMA8% and active agents such as a mix of growth factors or the antimicrobial enzyme lysozyme. The rolled tube is a stable construct that could serve as a neurotube, promoting the regeneration of peripheral nerves. Additionally, we developed 3D bioprinting of collagen-cell-laden hydrogels on both ends of the tubes to imitate the growth of DRG neurons after injury. This developed *ex vivo*-like model could serve as a platform to stimulate neural regeneration by studying DRG migration or, in the future, other cell types or their combinations. We effectively demonstrated possibilities to study neural cell behaviour on these bioprinted constructs with and without loaded growth factors, with the influence of PEMF over a prolonged amount of time, and even the possibility to incorporate antimicrobial agents to study anti-infective strategies. The unique composition of the fibrous tubular architecture, composite hydrogel, growth factor composition, and PEMF enhanced D11 DRG neuron migration, neurite outgrowth, and cell-cell interaction, supporting their maturation and proving similar effects observed in *vivo* studies by others.[34] We expect our G***rooveNeuroTube*** model to be broadly applicable for future neural tissue developmental and regeneration studies and demonstrate its potential as an effective new graft for peripheral nerve repair.

## Acknowledgements

We are grateful to Dr. Abhishek Indurkar (Center for Advanced Technologies, Adam Mickiewicz University) for performing FTIR measurements and for his valuable contribution to the analysis of FTIR, NMR spectra, and swelling test results.

## CRediT authorship contribution statement

**J.L.**: Conceptualization, data curation, formal analysis, funding acquisition, investigation, methodology, project administration, resources, supervision, validation, visualization, co-writing - original draft. **Y.R.:** Writing – original draft, Validation, Methodology, Formal analysis. **H.I.:** Writing, Methodology, Investigation, Data curation. **A.O.:** Writing, Methodology, Investigation, Data curation. **Ł.P.:** Writing, Methodology, Investigation, Data curation. **K.F.:** Writing, Methodology, Investigation, Data curation. **J.C.R.C.:** Writing, Methodology, Investigation, Data curation. **J.K.W.**:, Formal analysis, methodology, validation, visualization, co-writing - original draft; **K.T**.: Writing, Methodology, Investigation, visualization, Data curation.

## Notes

The authors declare no competing financial interest. All research data supporting this publication are directly available within this publication, as well as associated supporting information.

