## Supporting Information for "3D Bioprinted Cell-laden GrooveNeuroTube: A Multifunctional Platform for *Ex Vivo* Neural Cell Migration and Growth Studies"

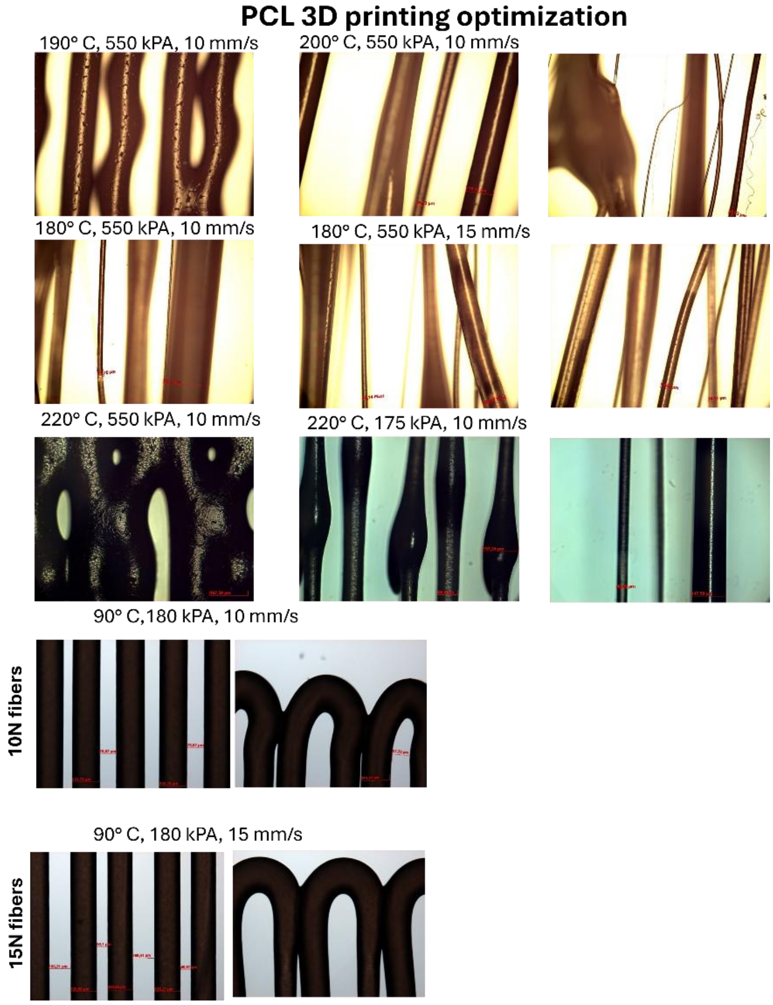


***Figure S1:*** *SEM Morphological characterization of PCL fibers optimization*

Table 1 - List of primers used for RT-PCR

| Gene | Primer | Accession Number | Primer Sequence (5' → 3') |
|---|---|---|---|
| **DCX** | Forward | NM_053379.3 | TCAGGTAACGACCAAGACGC |
| **DCX** | Reverse | NM_053379.3 | CAGGGCTTGTGGGTGTAGAG |
| **MAP2** | Forward | NM_013066.1 | CTTGATTCTATTGCCCTTGGGTTTA |
| **MAP2** | Reverse | NM_013066.1 | CATCCATCGTTCCGCTAGTGTTG |
| **Peripherin** | Forward | NM_012633.3 | ATCTCAGTGCCCGTTCATTC |
| **Peripherin** | Reverse | NM_012633.3 | AGCAGGACTGGTTGCAGACT |
| **NF200** | Forward | NM_012607.2 | GCAGTCAGAGGAGTGGTTCC |
| **NF200** | Reverse | NM_012607.2 | TCTCAATATCCAGGGCCATC |
| **SYP** | Forward | NM_012664.3 | CGGAATACTTGGAGGCTGGG |
| **SYP** | Reverse | NM_012664.3 | ACAATACCGAAGGGCACAGG |
| **SYN** | Forward | NM_019133.2 | CCAATGCCTTCAACCTTCCAG |
| **SYN** | Reverse | NM_019133.2 | GCGGATGGTCTCAGCTTTCA |
| **S100b** | Forward | NM_013191.2 | GGAGCTCATCAACAACGAGC |
| **S100b** | Reverse | NM_013191.2 | GGAAGTCACACTCCCCATCC |
| **TUBB3** | Forward | NM_139254.2 | CAACTATGTGGGGGACTCGG |
| **TUBB3** | Reverse | NM_139254.2 | TGGCTCTGGGCACATACTTG |
| **GAPDH** | Forward | NM_017008.4 | GGGCTGCCTTCTCTTGTGAC |
| **GAPDH** | Reverse | NM_017008.4 | TTTCCCGTTGATGACCAGCTT |
| **ACTB** | Forward | NM_031144.3 | GAAGATCCTGACCGAGCGTG |
| **ACTB** | Reverse | NM_031144.3 | GCTCGAAGTCTAGGGCAACA |

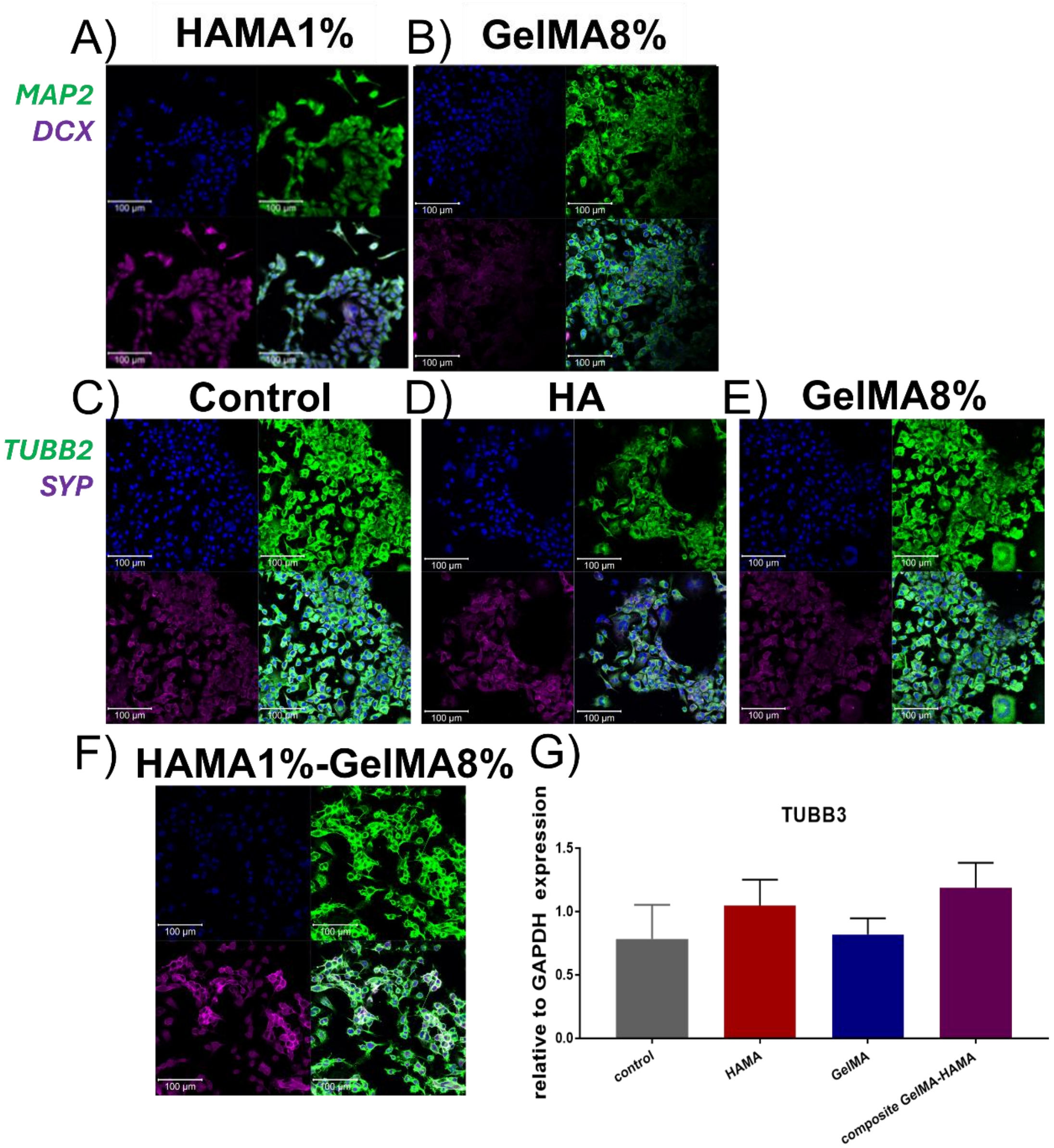


***Figure S2***. *ICC detection of migrating (DCX, violet) and late (MAP2, green) neuronal markers in cells cultured on A) only HAMA1% and B) only GelMA8% hydrogels for 14 days. ICC detection of synaptic vesicles (SYP, violet) and late (TUBB2, green) neuronal markers in cells cultured on C) control Petri Dish, B) only HAMA1%, C) only GelMA8%, F) composite HAMA1%-GelMA8% hydrogels for 14 days. G) The chart presents the differences in relative expression of the TUBB3 genes (relative to GAPDH) for F11 DRG neural cell line cultured on top of HAMA, GelMA and composite HAMA1%-GelMA8%. Statistical analysis was performed*

*using One Way Anova with post hoc Tukey's test. Data are presented as mean ± standard deviation; n=3, *p<0.05, **p<0.01, ***p<0.001, ****p<0.0001.*

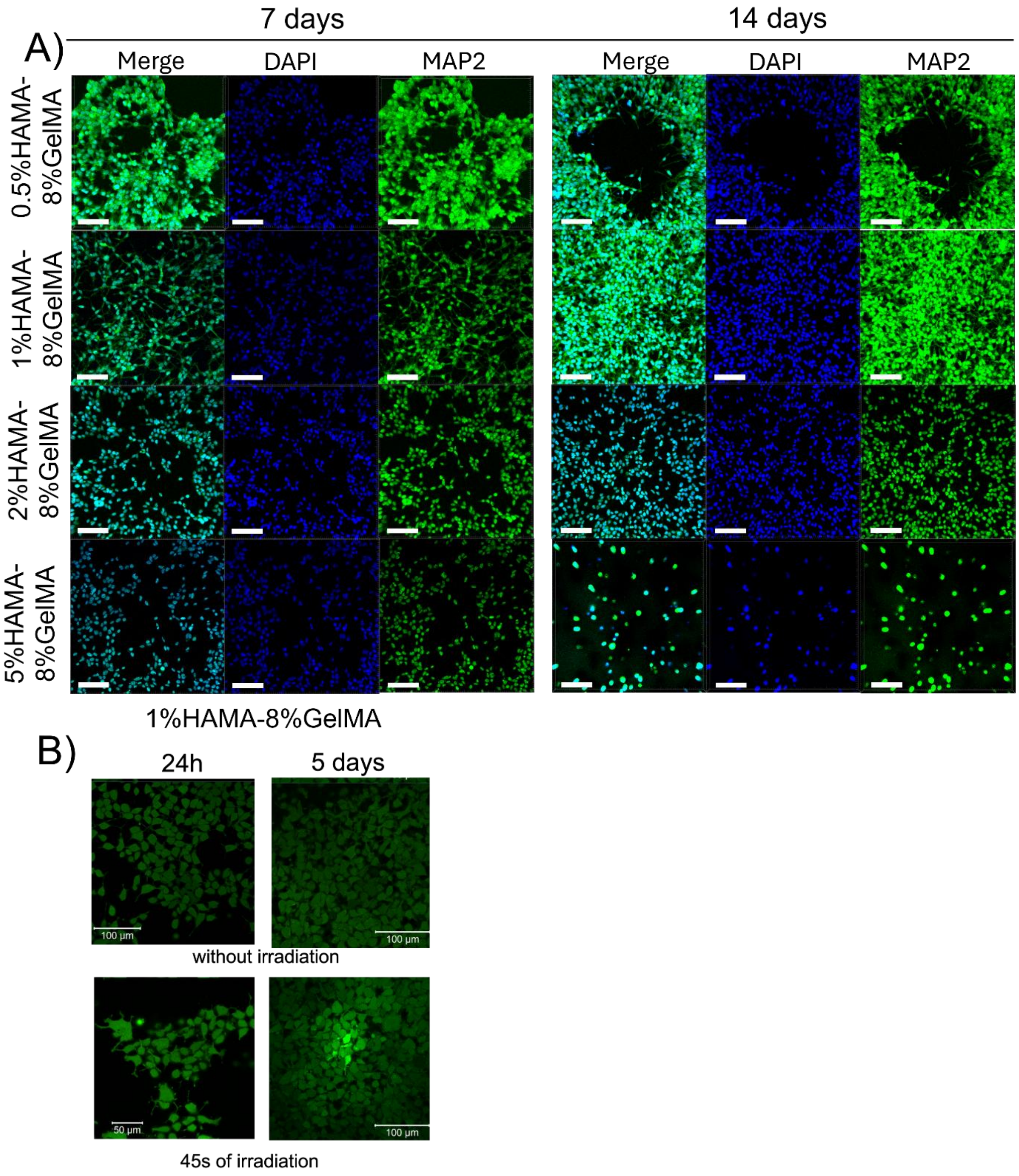

***Figure S3: A)*** *ICC detection of late (MAP2, green) neuronal marker in cells cultured in varying HAMA content (0.5%, 1%, 2%, and 5%)-8%GelMA for 7 and 14 days;* ***B)*** *Representative confocal images of live/cell assay of live (Calcein-AM labeled; green) and dead (EtDH-1, red) F11 DRG neural cells cultured on top of the hydrogel materials for 24 h and 5 days*

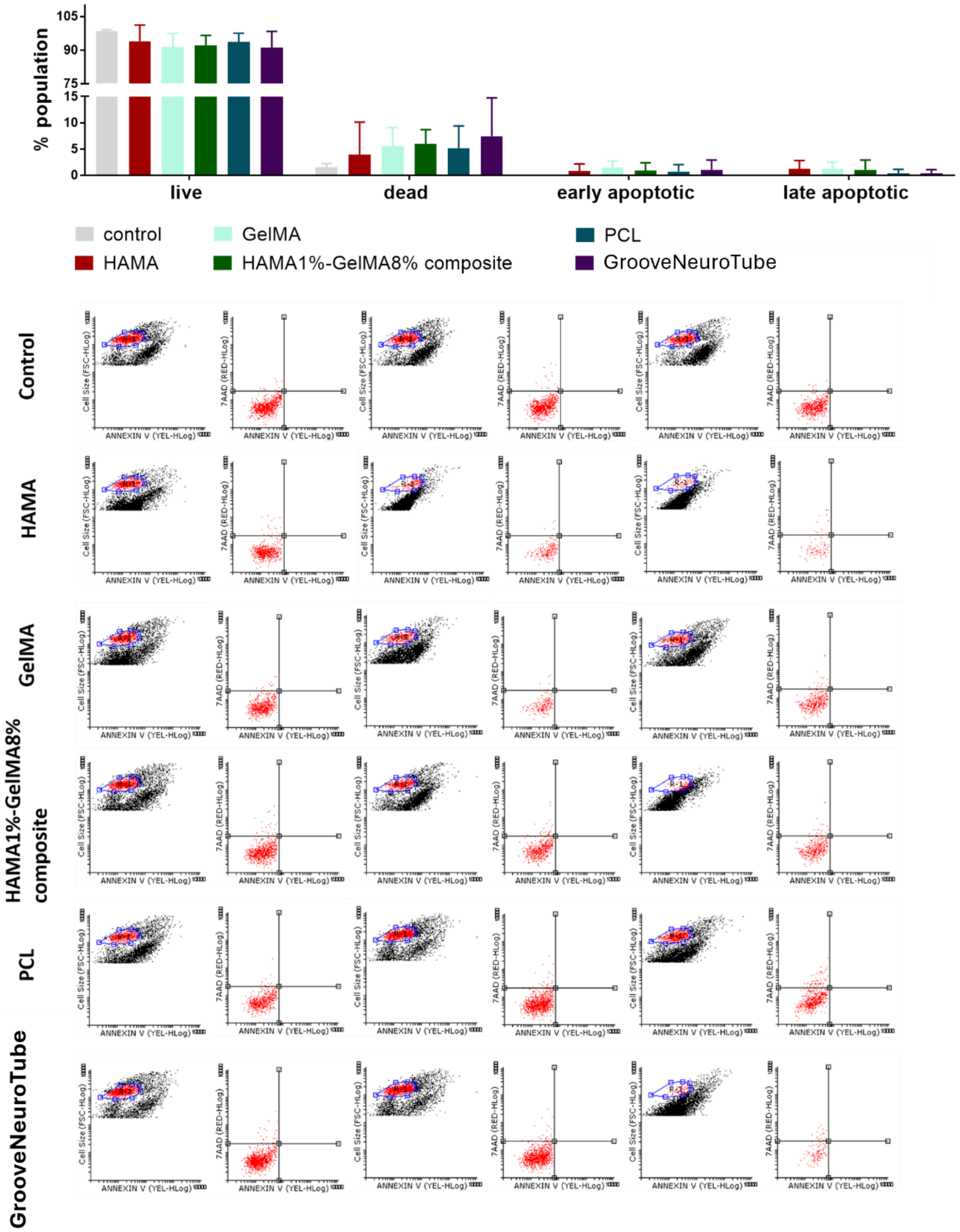


***Figure S4:*** *The viability test results for F11 DRG neural cells cultured on substrates made of hyaluronic acid methacrylate (HAMA), gelatin methacrylate (GelMA), a HAMA1%-GelMA8% composite, PCL, and a PCL-filled with HAMA1%-GelMA8%, were obtained. The experiment used the Muse™ Annexin V & Dead Cell Kit and was analyzed by flow cytometry. The*

*experiments were performed in three biological replicates to ensure data reliability and reproducibility.*

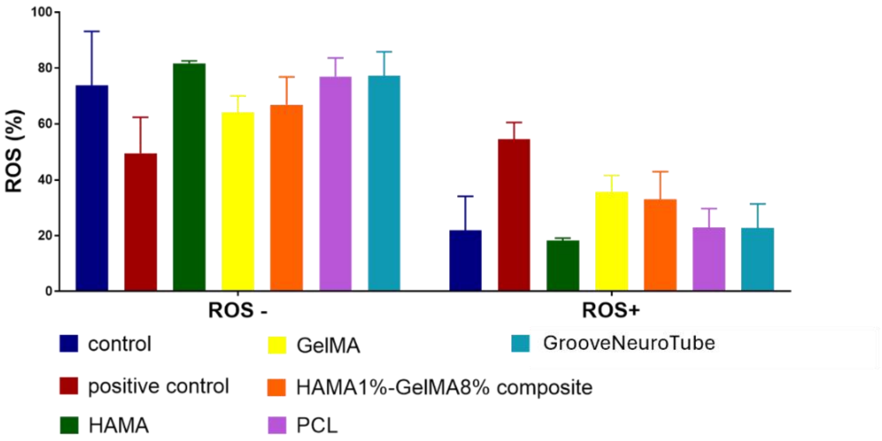
ROS (%)
100
80
60
40
20
0
ROS -
ROS+
control
positive control
HAMA
GelMA
HAMA1%-GelMA8% composite
PCL
GrooveNeuroTube

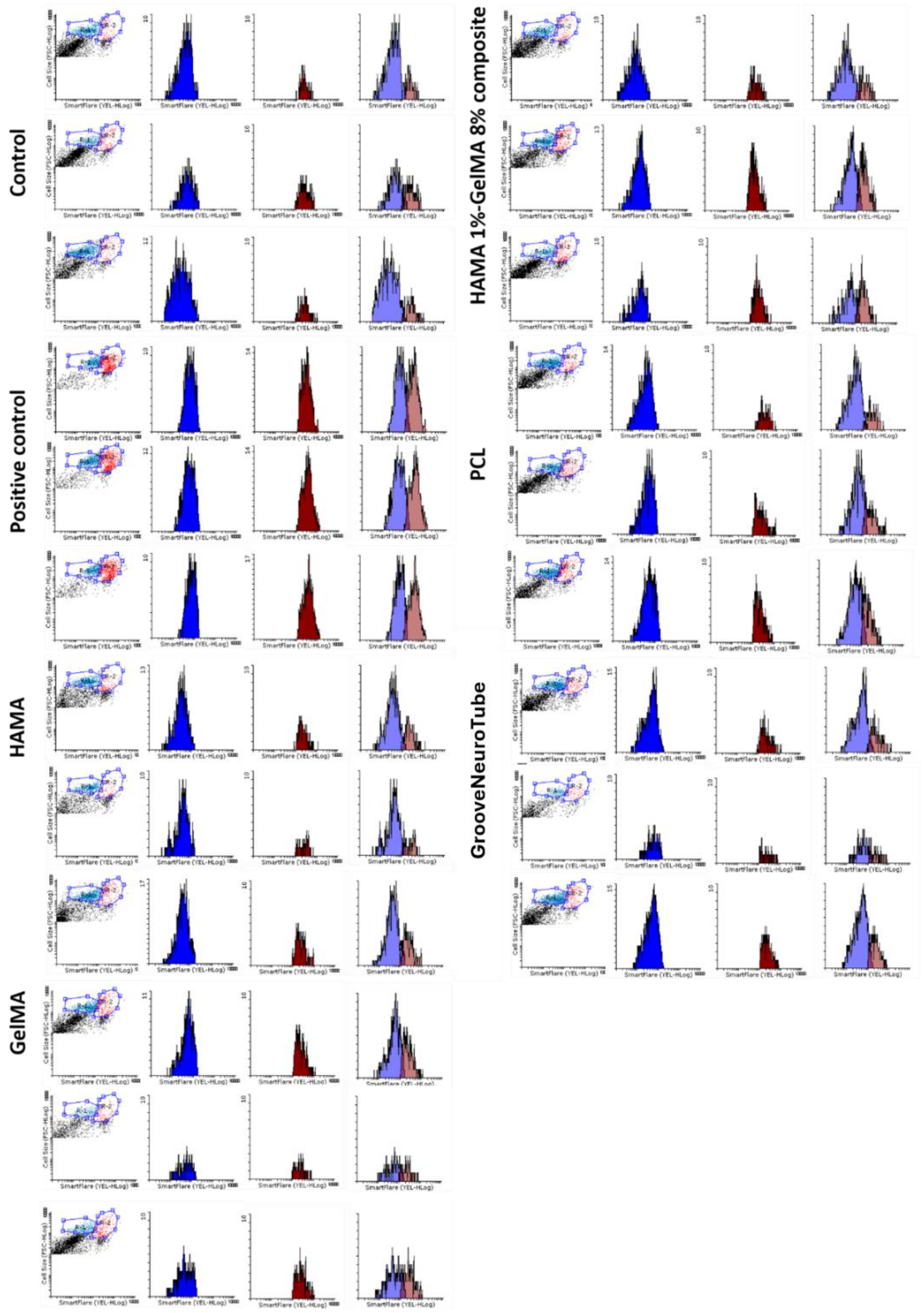
Control
Positive control
HAMA
GelMA
HAMA 1%-GelMA 8% composite
PCL
GrooveNeuroTube

***Figure S5:*** *Oxidative stress analysis: The generation of reactive oxygen species (ROS) in F11 DRG neural cells after 14 days of culture on various substrates was assessed. The R1 population, marked in blue, represents the ROS-negative (ROS-) population, while the ROS-positive (ROS+) population is marked in red. A summary graph illustrates the results of the oxidative stress test conducted on F11 DRG neural cells, highlighting the proportion of ROS+ and ROS- populations under the tested conditions.*

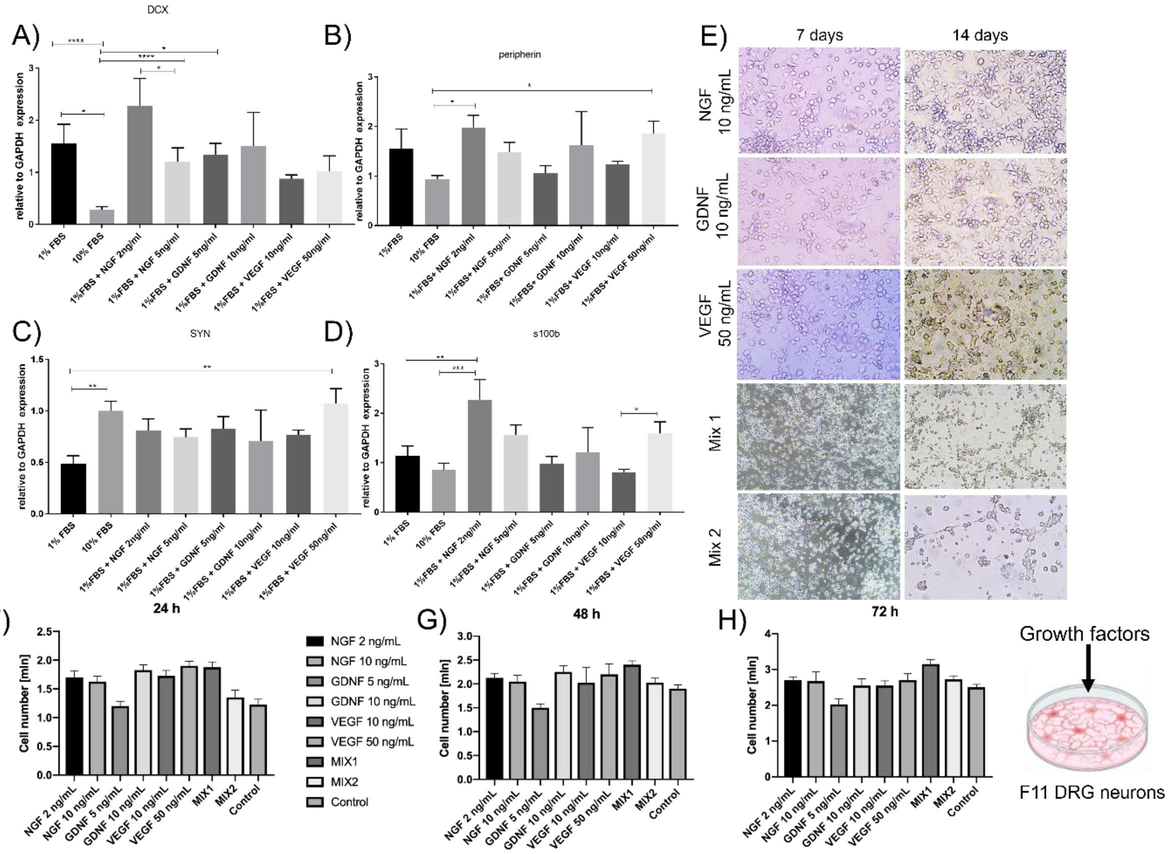


***Figure S6:*** *The differences in relative expression of the A) DCX, B) peripherin, C) SYN, D) s100b genes (relative to GAPDH) for F11 DRG neural cell line cultured in 1% FBS with the*

addition of growth factors (NGF 2 ng/mL and 5 ng/mL, GDNF 5 ng/mL and 10 ng/mL, as well as VEGF 10 ng/mL and 50 ng/mL). Statistical analysis was performed using One-Way ANOVA (performed in GraphPad software) with Tukey's post hoc test, n=3. Data are presented as mean ± standard deviation; *p<0.05, **p<0.01, ***p<0.001, ****p<0.0001. E) photos from an optical microscope showing the morphology of F11 cell cultures with growth factors for 7 and 14 days. F) Proliferation of F11 cells treated with growth factors for 1, 2, and 3 days.

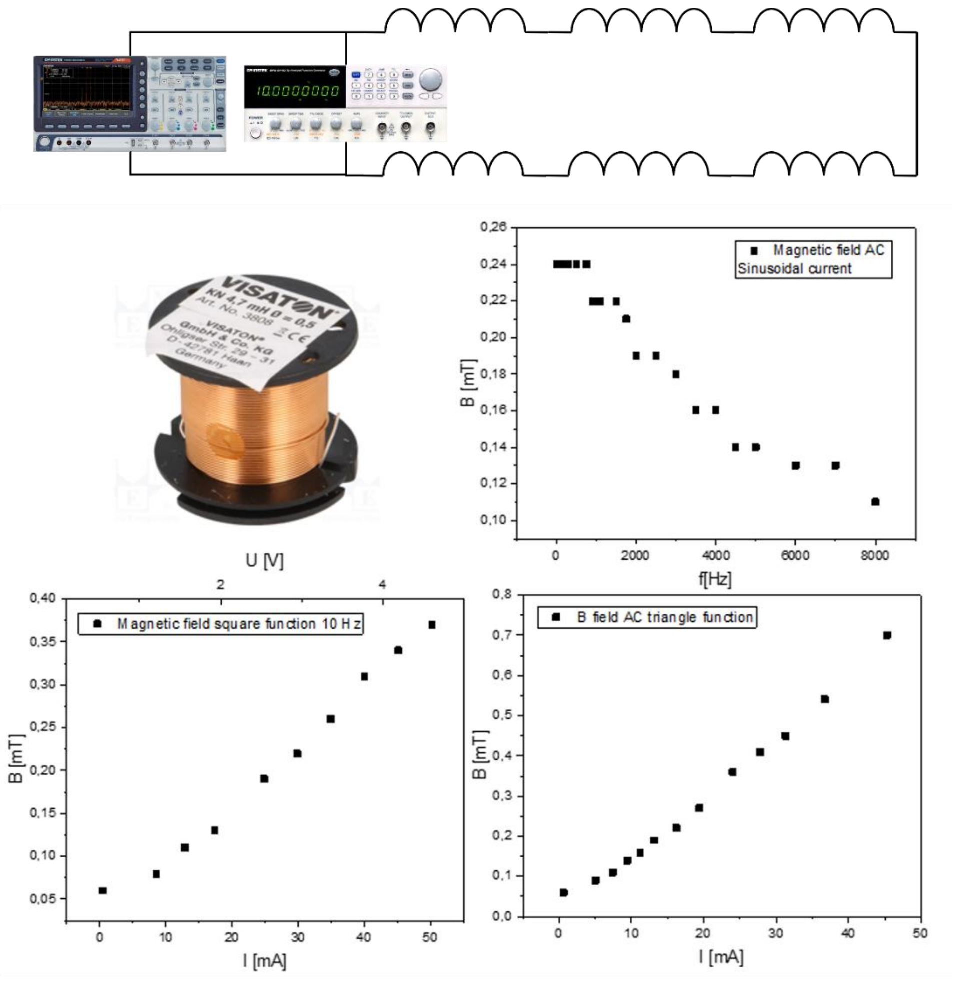


***Figure S7:*** *Scheme of the PEMF setup. Image of the coils used in that study and technical characterization of the magnetic fields from the coils: triangle, sinusoidal, and square waveform. An alternating current, e.g., sine, square, or triangle-shaped, induces a sinusoidally varying magnetic field.*

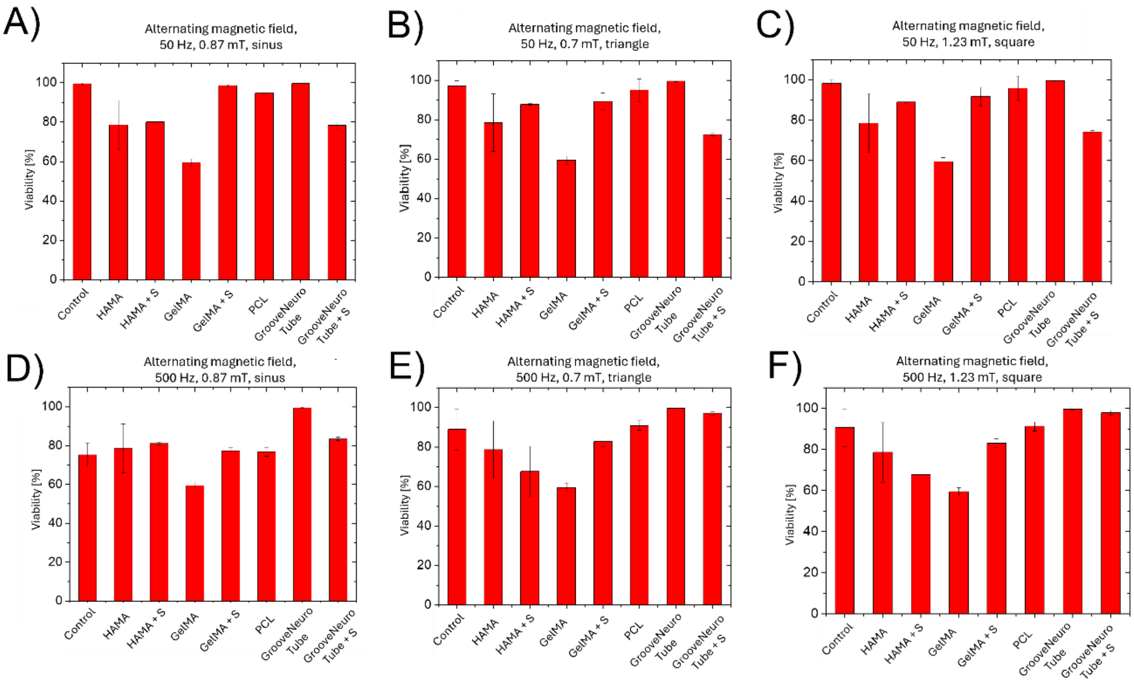


***Figure S8:*** *Viability of F11 DRG neural cells and human neural cells without any treatment (control) and treated with variable magnetic fields (A, D sinus; B, E triangle and C, F square) with a frequency of A, B, C) 50 Hz (sinus waveform) or D, E, F) 500 Hz (square waveform).*

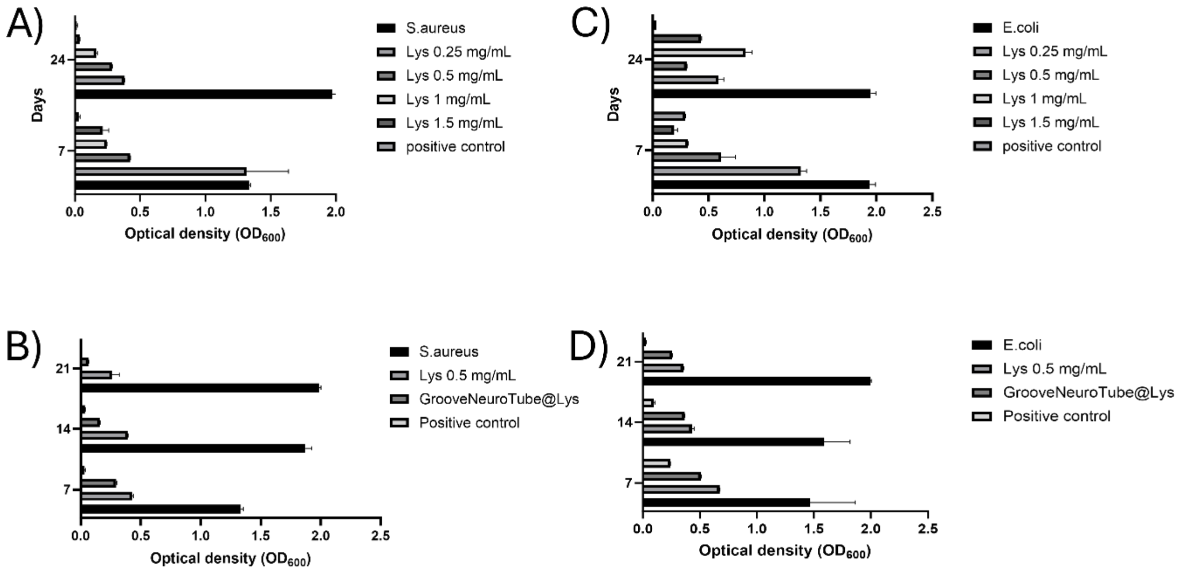

***Figure S9:*** *Antibacterial activity of lysozyme at a concentration range of 0.25-1.5mg/mL against E.coli and S.aureus, B) Antibacterial activity of crosslinked hydrogels, pristine PCL and lysozyme against E.coli, C) Antibacterial activity of crosslinked composite hydrogels, pristine PCL and lysozyme against E.coli after 7, 14 and 21 days in hydrogels incubation*

### Extended discussion on antimicrobial testing

We studied its effects over a more extended period (up to 21 days) against *S. aureus* and *E. coli*. We loaded lysozyme within HAMA1%-GelMA8% hydrogel and studied inhibition bacteria over 7, 14, and 21 days. As expected, freely loaded lysozyme is more effective in inhibiting *E. coli* and *S. aureus* than lysozyme encapsulated within the HAMA1%-GelMA8% hydrogel. However, it is still sufficiently effective in inhibiting over 80% of the bacteria within 21 days (***Figure S9***). Different concentrations of lysozyme were also tested with F11 DRG neural cells; no cytotoxic effects, reduction in proliferation, or changes in morphology were observed (***Figure S10***).

After 7, 14, and 21 days of incubation, *S. aureus* was exposed to PBS supernatant containing lysozyme released from the GelMA-HAMA hydrogel, and OD600 measurements were taken after 24 hours. On day 7, the OD600 for *S. aureus* in the GelMA-HAMA-Lyz group was 0.295, much lower than the positive control (1.335), indicating effective antibacterial action. By day 14, the OD600 decreased to 0.159, while the positive control remained high (1.872). On day 21, the GelMA-HAMA-Lyz group exhibited the most substantial effect with an OD600 of 0.068, compared to the positive control (1.987), showing sustained antibacterial efficacy over time. The lysozyme solution alone (Lyz 0.5 mg/mL) showed moderate antibacterial effects but was less effective than the hydrogel. These results highlight the hydrogel's ability to maintain antibacterial activity over an extended period (***Figure S9B***). The GelMA-HAMA-Lyz hydrogel showed significant antibacterial activity against E. coli. On day 7, the OD600 for E. coli in the GelMA-HAMA-Lyz group was 0.509, lower than the positive control (1.466). By day 14, the OD600 decreased further to 0.369, while the positive control remained high at 1.592. On day 21, the GelMA-HAMA-Lyz group showed the most significant reduction in bacterial growth, with an OD600 of 0.257, compared to the positive control (1.994). The lysozyme solution alone (Lyz 0.5

mg/mL) demonstrated moderate antibacterial effects, but the GelMA-HAMA-Lyz hydrogel was more effective in reducing bacterial growth over time. These results indicate the sustained antibacterial potential of the GelMA-HAMA-Lyz hydrogel against E. coli (***Figure S9D***). This confirmed that the ***GrooveNeuroTube*** construct can be used to study the effects of antibacterial agents to mimic infection-like scenarios in future studies.

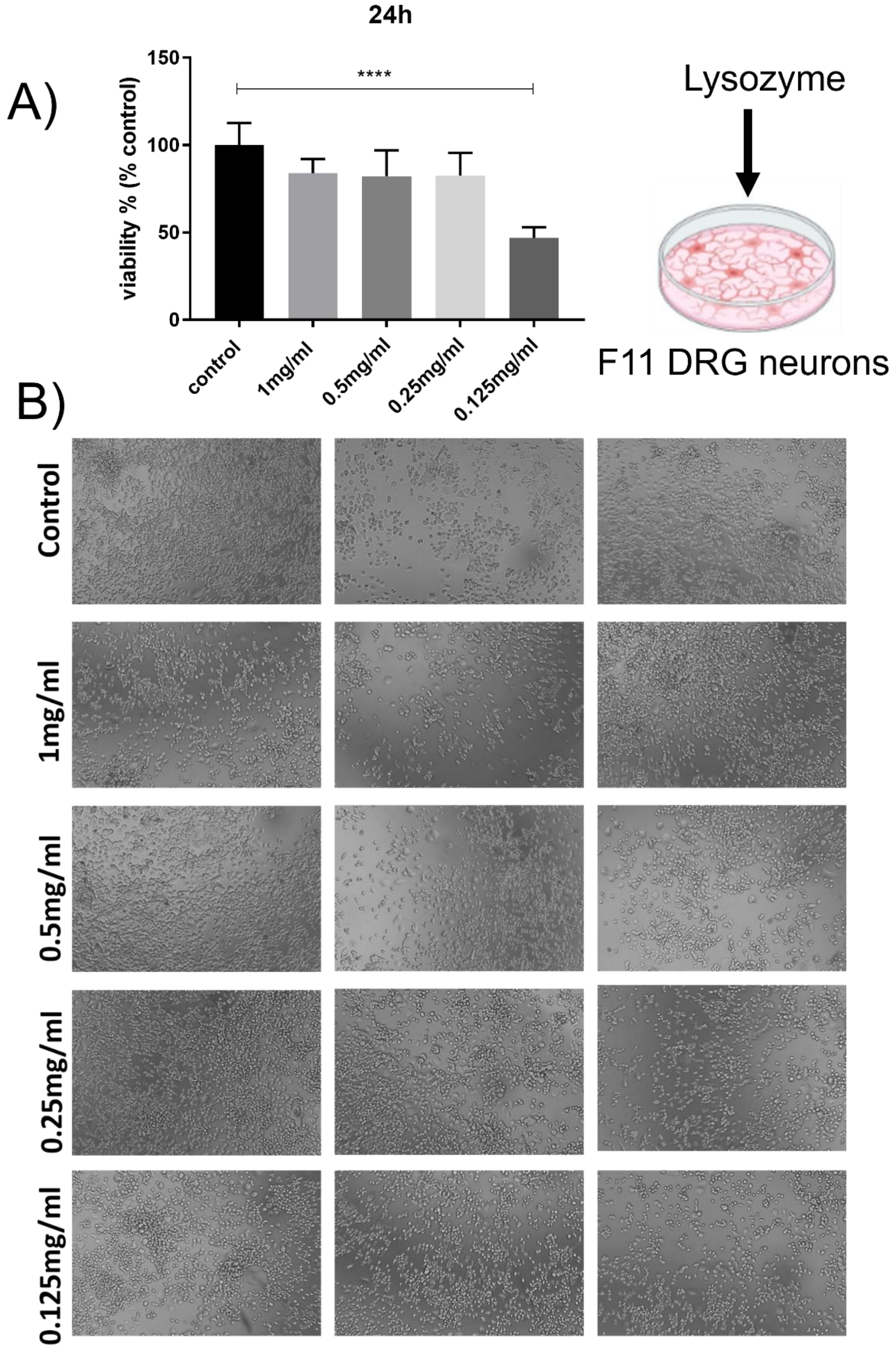


***Figure S10:*** *Effect of lysozyme treatment concentration for 24 hours on F11 DRG neural cells: A) viability and B) morphology*

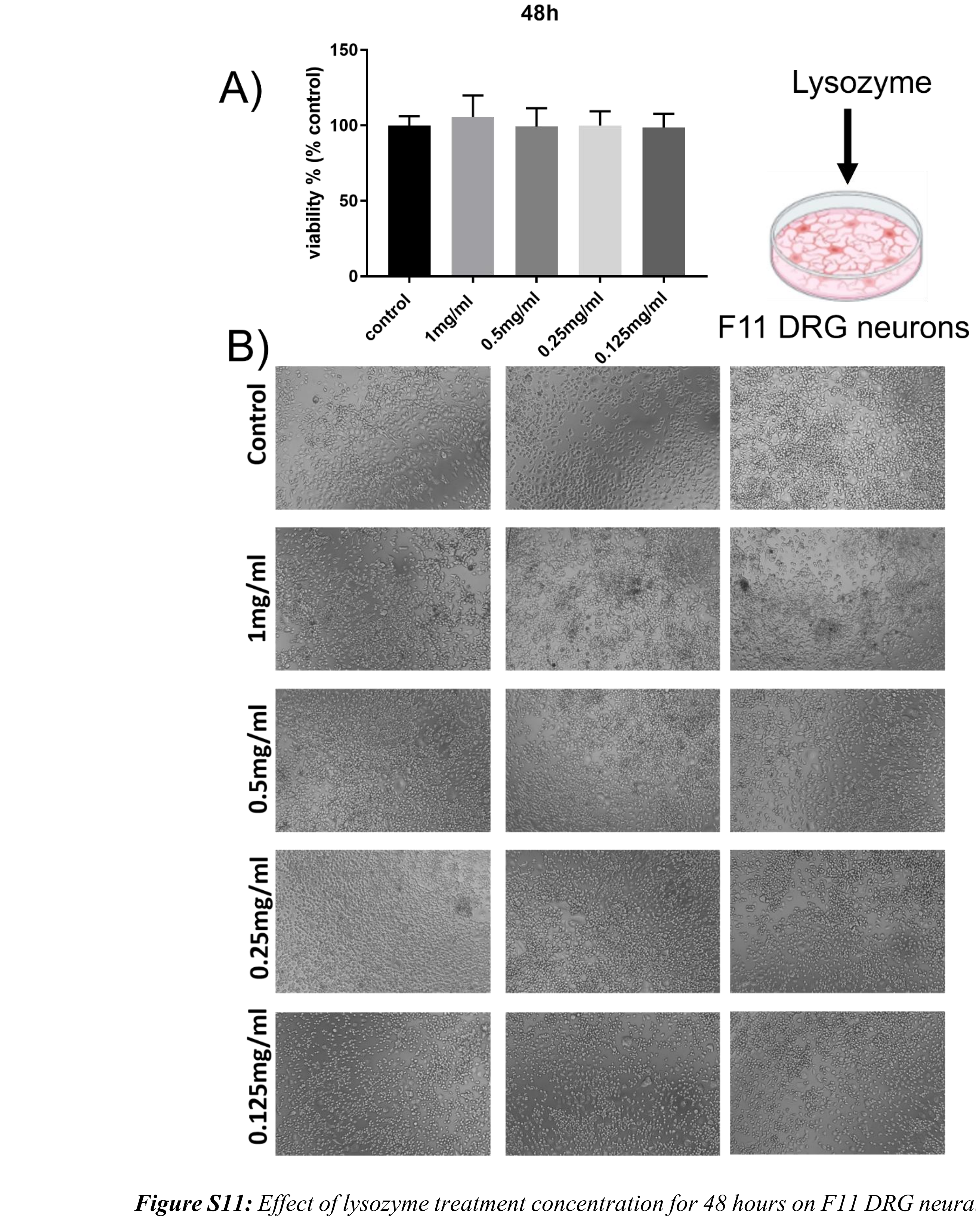


***Figure S11:*** *Effect of lysozyme treatment concentration for 48 hours on F11 DRG neural cells: A) viability and B) morphology*

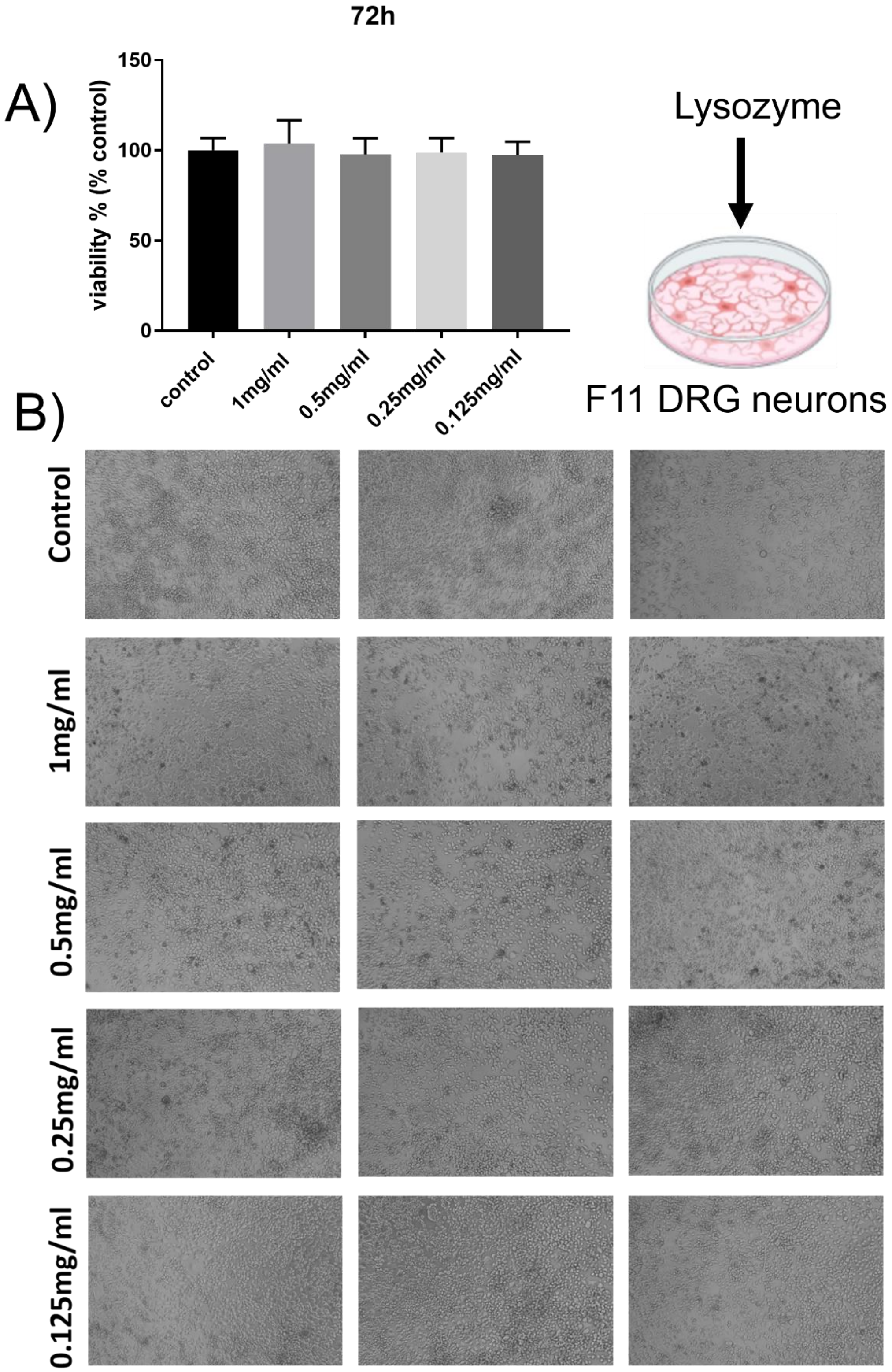


***Figure S12:*** *Effect of lysozyme treatment concentration for 72 hours on F11 DRG neural cells: A) viability and B) morphology*
